# Contrasting recruitment of skin-associated adipose depots during cold challenge of mouse and human

**DOI:** 10.1101/2020.09.16.300533

**Authors:** Ildiko Kasza, Jens-Peter Kühn, Henry Völzke, Diego Hernando, Yaohui G. Xu, John W. Siebert, Angela LF Gibson, C.-L. Eric Yen, David W. Nelson, Ormond A. MacDougald, Nicole E. Richardson, Dudley W. Lamming, Philip A. Kern, CM Alexander

## Abstract

Mammalian skin impacts metabolic efficiency system-wide, controlling the rate of heat loss and consequent heat production. Here we compare the unique fat depots associated with mouse and human skin, to determine whether they have corresponding function and regulation. For human, we assay a skin-associated fat (SAF) body-wide depot to distinguish it from the subcutaneous fat pads characteristic of abdomen and upper limbs. We show that the thickness of SAF is not related to general adiposity; it is much thicker (1.6-fold) in women than men, and highly subject-specific. We used molecular and cellular assays of β-adrenergic induced lipolysis and found that dermal white adipose tissue (dWAT) in mice is resistant to lipolysis; in contrast, the body-wide human SAF depot becomes lipolytic, generating heat in response to β-adrenergic stimulation. In mice challenged to make more heat to maintain body temperature (either environmentally or genetically), there is a compensatory increase in thickness of dWAT: A corresponding β-adrenergic stimulation of human skin adipose (*in vivo* or in explant) depletes adipocyte lipid content. We summarize the regulation of skin-associated adipocytes by age, sex, and adiposity, for both species. We conclude that the body-wide dWAT depot of mice shows unique regulation that enables it to be deployed for heat preservation; combined with the actively lipolytic subcutaneous mammary fat pads they enable thermal defense. The adipose tissue that covers human subjects produces heat directly, providing an alternative to the brown adipose tissues.

**KEY POINTS SUMMARY:**

- Several distinct strategies produce and conserve heat to maintain body temperature of mammals, each associated with unique physiologies, with consequence for wellness and disease susceptibility
- Highly regulated properties of skin offset the total requirement for heat production
- We hypothesize that the adipose component of skin is primarily responsible for modulating heat flux; here we evaluate the relative regulation of adipose depots in mouse and human, to test their recruitment to heat production and conservation
- We found that insulating mouse dermal white adipose tissue accumulates in response to environmentally- and genetically-induced cool stress; this layer is one of two adipose depots closely apposed to mouse skin, where the subcutaneous mammary gland fat pads are actively recruited to heat production
- In contrast, the body-wide adipose depot associated with human skin produces heat directly, potentially creating an alternative to the centrally regulated brown adipose tissue

## INTRODUCTION

Specializations of mammalian adipose depots determine their individual impact on human health (Manolopoulos *et al*., 2010; Karastergiou & Fried, 2013; Lee *et al*., 2013; Yoneshiro *et al*., 2013; Pinnick *et al*., 2014; Walker *et al*., 2014). In general, obesity is associated with metabolic abnormalities and the development of a variety of diseases, from diabetes to cancer, yet not all large deposits of adipose tissue types are linked to disease susceptibility (Smith *et al*., 2019). For example, intraperitoneal visceral adipose tissue (vWAT) is linked to cardiometabolic risk, whereas accumulation of subcutaneous white adipose tissue (scWAT), especially gluteofemoral adipose in the lower body, appears to protect subjects from these health risks (Manolopoulos *et al*., 2010; Karastergiou *et al*., 2012).

Studies of scWAT typically describe the gluteofemoral and/or abdominal depots; however, even high-resolution techniques such as magnetic resonance imaging (MRI) tend to overlook the body-wide fat layer underneath with skin. Our studies have drawn attention to this depot; indeed a preliminary calculation suggested that this body-wide depot comprised at least half of the fat for the average lean woman (Kasza *et al*., 2016).

The term “subcutaneous white adipose tissue” is an umbrella term that has been used differently by previous studies, and is likely to include a heterogeneous set of depots and adipocyte cell types (Kelley *et al*., 2000; Driskell *et al*., 2014; Walker *et al*., 2014; Kruglikov & Scherer, 2016b; Nicu *et al*., 2018; Zwick *et al*., 2018). For this study we therefore distinguish a body-wide depot (**skin-associated fat, SAF)** from **subcutaneous fat pads (sc fat pads; Fig. 1A)**. SC fat pads accumulate around the abdomen and upper limbs, particularly in over-weight women with gynoid (pear-shaped) fat deposition (Wajchenberg, 2000; Karastergiou & Fried, 2017), and they have been quantified by MRI (Smith *et al*., 2001). Our goal is to specifically examine the behavior, properties and regulation of skin-associated fat as it relates to heat transfer; therefore, given its size and location, we quantified SAF.

**Fig. 1.**
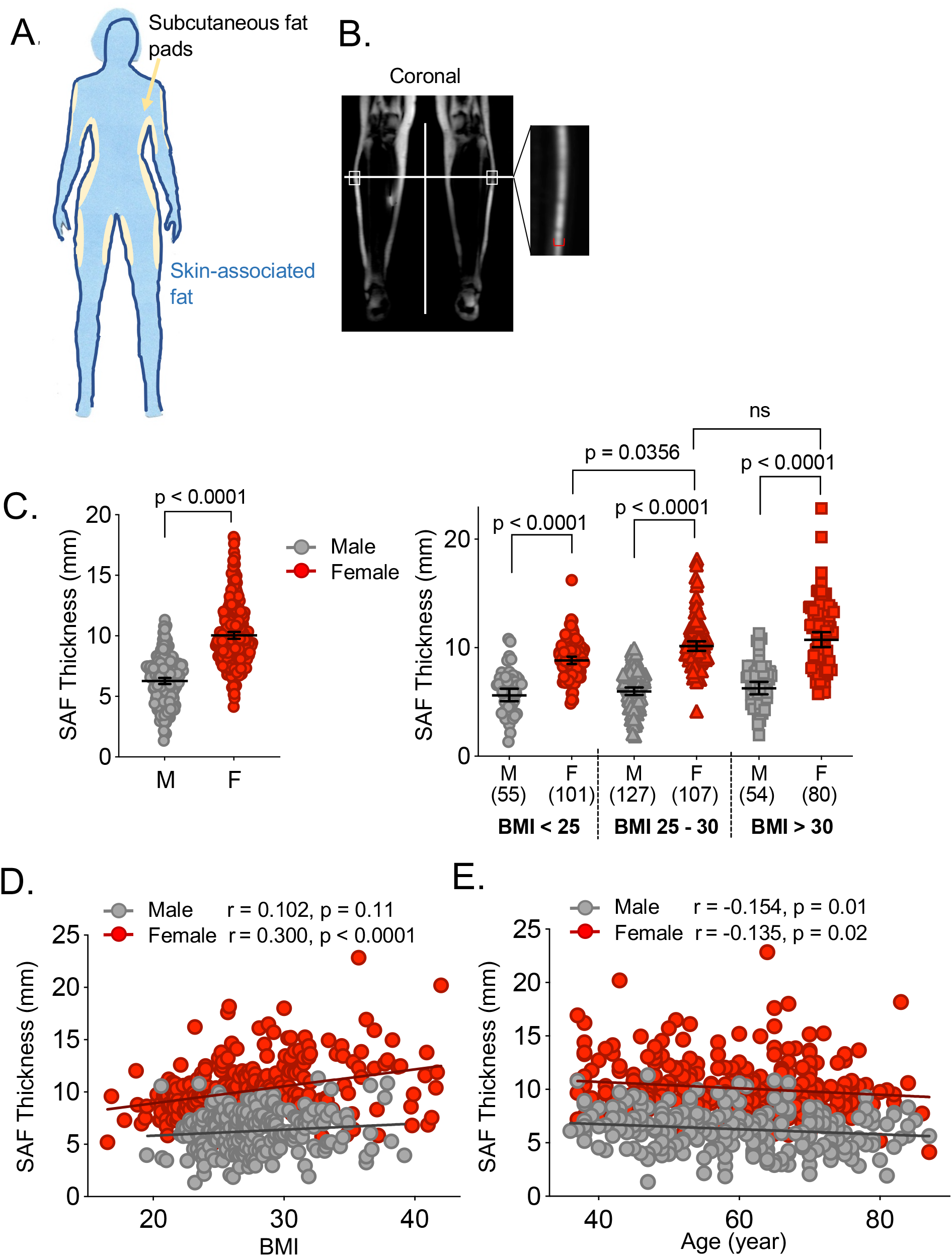
The thickness of skin-associated fat is highly variable between human subjects, and independent of BMI. **A.** Cartoon illustrates the use of assay terminology. Ectopic subcutaneous fat pads are colored yellow; the body wide depot of skin associated fat colored dark blue. **B**. *Assay of skin associated fat* measured using coronal, fat-only MRI views of lower leg, 30% of distance between knee and ankle. **C**. Results of SAF thickness assay for men (n=236) and women (n=286) as determined by MRI: these data are shown together (left hand side,) and after dividing the subjects into groups, namely lean (BMI<25), overweight (BMI 25-30) and obese (BMI>30), with the indicated groups sizes (right hand side). Data are represented as a geometric mean with 95% confidence interval. The significance of differences was analyzed by Kruskal-Wallis tests corrected for multiple comparisons with a Dunn’s test. **D, E**. Evaluation of correlation between BMI, age and SAF thickness, for men and women, indicated using Pearson’s correlation coefficients.

We are focused on the thermal defense properties of mouse and human skin-associated fat. Thus, a layer of fat, applied body-wide to skin, is predicted to impact heat transfer properties dramatically. In turn, this affects the metabolic budget and energy expenditure of mammals; for example, mice housed at temperatures 10^0^C less than their thermoneutral temperature show a 50% increase in energy expenditure (Tschop *et al*., 2012; Speakman, 2013; Kasza *et al*., 2014). A thermoneutral environment for mice is warmer than for humans (approximately 31°C compared to room temperature/24°C), and the effect of room temperature housing on the physiology of mice is considerable, suppressing a wide range of inflammatory reactions (Tian *et al*., 2016; Giles *et al*., 2017; Qiao *et al*., 2018).

The dramatic physiological cost of heat production has not gone unnoticed; a major research effort has been directed to devising a strategy for treating the epidemic rates of obesity in westernized populations, by stimulating thermogenic adipose depots (beige/brown) to “burn” calories, using either direct cold exposure or β-adrenergic agonists (Cannon & Nedergaard, 2009; Harms & Seale, 2013; Yoneshiro *et al*., 2013; Bartelt & Heeren, 2014; Elattar & Satyanarayana, 2015). The physiology that results from beige adipose activation is different from that arising from brown adipose depots (Keipert & Jastroch, 2014; Keipert *et al*., 2020), since each adipose depot is endocrine and highly specialized (Kershaw & Flier, 2004; Villarroya & Giralt, 2015; Wang *et al*., 2015). It is not known how the relative contribution of each mechanism is determined.

Our previous studies have shown that the skin-associated fat depot of mice (dermal white adipose tissue, dWAT) is highly regulated in a manner that suggests that this depot is part of an acclimation process that mitigates heat loss (Alexander *et al*., 2015). Thus dWAT thickens in mice housed at room temperature (sub-thermoneutral), and mice that are unable to develop thicker dWAT are chronically cold stressed (Kasza *et al*., 2014). Importantly, several mouse strains with defects of skin lipogenic enzymes lose heat at high rates; interestingly, these strains are also resistant to diet-induced obesity (Sampath *et al*., 2009; Shih *et al*., 2009; Neess *et al*., 2013; Sampath & Ntambi, 2014; Kruse *et al*., 2017; Kasza *et al*., 2019; Neess *et al*., 2020).

Here, we aim to provide insight on the functional homologies between the skin-associated adipose depots in mouse and human, so that models aimed at testing the role of the regulated properties of skin in systemic metabolism can be built more accurately. We quantify not just skin thickness, but skin-associated fat, for human subjects, and find that it is surprisingly variable between individuals. We show that this fat can directly produce heat, suggesting a highly tailored strategy for maintaining body temperature that varies between individuals depending upon the size of their skin-associated fat depots. We derive data from a genetically cool-stressed mouse model that shows that the accumulation of dWAT is a component of the UCP1-independent heat generating response, which together with a demonstration that this depot is entirely insensitive to β-adrenergic agonists, supports our claim that dWAT is designed as a functional insulator within mouse skin.

Overall, we show that the mouse skin-associated adipose depots comprise two distinct types of fat, one insulating (dWAT), and the other a lipolytic thermogenic depot (scWAT). In contrast, the human skin-associated adipose tissues are surprisingly homogeneous with respect to their responsivity to thermogenic cues, regardless of depth from skin, or body site; indeed we could find no evidence for extensive colonization of human skin by a mouse dWAT equivalent.

## METHODS

### Ethical Approval. Mice

These studies were performed in strict accordance with the recommendations in the Guide for the Care and Use of Laboratory Animals of the National Institutes of Health. Experimental protocols were approved by the University of Wisconsin School of Medicine and Public Health Animal Care and Use Committee. The number of mice used to perform this study was minimized, and every effort was made to reduce the chance of pain or suffering. Method of euthanasia was CO2 asphyxiation, as per guidelines. All authors understand the ethical principles operating at the Journal of Physiology and confirm that this work complies with the animal ethics checklist.

### Human Subjects

Subjects submitted their informed consent for the SHIP study as part of the Community Medicine Research Net of the University of Greifswald, Germany; the University of Greifswald is a member of the ‘Center of Knowledge Interchange’ program of Siemens AG, and this study was compliant with the Declaration of Helsinki. The Study of Health in Pomerania (SHIP) study is a longitudinal study of health parameters for over 4000 adults (20-79), representative of an area with population >200 000 (Volzke *et al*., 2011). All analysis of the SHIP data analyzed here was HIPAA-compliant and performed after obtaining approval from the ethics committee of the University of Griefswald (Germany) and the University of Wisconsin Institutional Review Board. Human skin specimens were taken from discarded tissues during defect reconstruction and body sculpting procedures and submitted, anonymously, to the BioCore Biobank, with IRB approval.

### Mice

BALB/cJ (#00651), C57BL/6J (#00664), Agouti lethal yellow (A^y^/a; C57BL/6J;#002468) and *UCP1-/-* C57BL/6J;#003124) mice were obtained from Jackson Labs and bred in-house. All animals were housed at constant temperature (19-23^0^C) in 12 h light/dark cycles with free access to water and standard chow (Harlan Teklad Global Diet 2020X). For high fat feeding, C57BL/6J mice were provided with Harlan Teklad diet#TD06415 (45% calories from fat); for calorie restriction (**CR**), *ad libitum* consumption rates of chow were measured for singly housed mice; mice were 25% calorie restricted using a 60% mix of AIN93M:40% CR (Bioserv cat#F05314) in control diet (AIN93M, Bioserv cat#F05312), pair-fed daily at 5 pm, for 5 weeks.

### Histological Analysis

Skin, BAT, perigonadal WAT (vWAT) and mammary gland (iWAT) were dissected for histological processing as follows: Samples were paraformaldehyde-fixed (4%) overnight / 4^0^C and then paraffin-embedded for evaluation. Tissue sections were deparaffinized, re-hydrated and stained with either standard H&E or Trichrome protocols for visualization. For assay of dWAT thickness, 6 images of H&E-stained, non-anagen fields of skins (equivalent to ≥4500 μm linear dWAT) were assayed by image analysis (dividing total area by length). For immunofluorescent analysis, tissues were processed for heat-induced epitope retrieval, epitopes were blocked in 10% goat serum for 3 hours; samples were either incubated with Alexa-conjugated primary antibodies for 1 hour, or incubated overnight with the primary antibody, washed and incubated for 1h with secondary antibodies. Samples were visualized on a confocal microscope (Nikon A1RS Confocal Microscope). For fluorescent intensity assessment, at least 3 independent fields were obtained for each mouse sample, and signal was quantified using open source Fiji image processing package. In each case, the specific signal was normalized to the signal from lipid droplet–associated perilipin. For quantification of lipid droplet size, six independent fields were obtained and quantified using open source Fiji image processing package (https://loci.wisc.edu/software/fiji). Antibodies and immunohistochemical reagents were as follows: anti-CD31 (#3528; RRID:AB_2160882), anti-CD31 (#77699; RRID:AB_2722705), anti-Fabp4 (#3544; RRID:AB_2278527), anti-Fatty acid synthase (FASN; #3180; RRID:AB_2100796), anti-HSL (#4107; RRID:AB_2296900), anti-pS565 HSL (#4137; RRID:AB_2135498), anti-pS660 HSL (#4126; RRID:AB_490997), anti-pS 235/226 S6 (#2211; RRID:AB_331679), all from Cell Signaling Technology; Alexa Fluor488 anti-perilipin (#NB110-40760AF488; RRID:AB_2167264) and AlexaFluor647 anti-perilipin (#NB110-40760AF647; RRID:AB_1849889) both from Novus Biologicals; anti-UCP1 (#ab10983, RRID:AB_2241462; Abcam); secondary reagents were AlexaFluor546 goat anti-rabbit (#A-11035; RRID:AB_143051), AlexaFluor546 goat anti-mouse (#A-11030; RRID:AB_144695) and AlexaFluor633 wheat germ agglutinin (#W21404) from Thermo Fisher Scientific. Specificity of antisera was confirmed by immunohistochemical evaluation of accredited scenarios (provided as Supplemental data).

### 3T3L1 cell culture

Mouse 3T3-L1 preadipocytes were from the American Tissue Culture Collection (ATCC; RRID: CVCL_0123) and were maintained in Dulbecco’s modified Eagle’s medium supplemented with 4.5 g/l of glucose (Life Technologies), 10% fetal bovine serum and 100-U/ml penicillin and streptomycin. Briefly, cells were differentiated into adipocytes using MDI medium (100 mg/ml 3-isobutyl-1-methylxanthine, 100 ng/ml dexamethasone and 1 mg/ml insulin, both from Sigma) for 4 days, followed by 1 mg/ml insulin for an additional 4 days (Kasza *et al*., 2014), and fixed for immunofluorescent staining using 3% paraformaldehyde (20 mins/4^0^C).

### β-adrenergic induction

The pan β-adrenergic agonist, isoproterenol hydrochloride (**ISO**) was from Sigma; the β3-adrenoreceptor agonists, CL 316,243 disodium salt (**CL**) was obtained from Tocris Bioscience (UK) and mirabegron (**Mira**) was from Cayman Chemical. For CL administration *in vivo*, mice were acclimated to thermoneutrality for 3 days, injected with 1 mg/kg CL, and euthanized for tissue collection 60 minutes later. For β-adrenergic stimulation *in vitro*, 3T3-L1 cells were treated with ISO (10 μM) for 40 minutes at 37^0^C. For β-adrenergic stimulation *ex vivo*, human fat samples (approx. 0.5cm in size) were treated with ISO (10 μM) for 40 minutes at 37^0^C. For β-adrenergic stimulation *in vivo*, human subjects were administered Mira for 10 weeks (50 mg/day), with abdominal adipose biopsies before and after treatment, as described previously (Finlin *et al*., 2018).

### FLIR imaging

To measure surface temperatures by infrared thermography, we used a hand-held FLIR T360 camera (FLIR Systems, Oregon). Pin drops in the software of the FLIR camera were used to record surface temperatures and quantified using FLIR Tools Advanced Thermal Analysis and Reporting software; each photograph is internally and externally calibrated to show actual temperatures (Kasza *et al*., 2019).

### SHIP study design and acquisition of MRI images of skin-associated fat

The Study of Health in Pomerania (SHIP) study is a longitudinal study of health parameters for over 4000 adults (20-79), representative of an area with population >200 000 (Volzke *et al*., 2011). The assay of the thickness of skin-associated fat (SAF) was performed using a chemical shift-encoded 3D gradient-echo sequence of the whole-body (1.5T MR images, Avanto, Siemens Healthcare), with the following parameters: repetition time = 12 ms, echo time (s) = 2.38 ms, flip angle = 5^0^, spatial resolution 1.95 mm x 1.95 mm x 5 mm, acquisition matrix 0/256/128/0, pixel bandwidth 1955 Hz/Pixel of each series with complete imaging of both calves. MR sequences were postprocessed to separate confounder corrected water- and fat-only images. Confounder-corrected fat only images were used to measure skin-associated fat (Kuhn *et al*., 2017). Skin associated fat was measured at the lateral side of both legs, 30% distance between tibia and ankle diaphysis.

### Collection of human samples

Breast tissues were collected during breast reduction surgeries by the University of Wisconsin Carbone Cancer center Translational Science Biocore Biobank. Human skin samples (with associated fat) were obtained from patients undergoing elective reconstructive surgeries at our institution. The de-identified samples were exempt from the regulation of University of Wisconsin-Madison Human Subjects Committee Institutional Review Boards. Data on patient age, sex, and type of surgery were collected with the tissues. Skin samples were processed within 3 hours of surgery.

### Statistical Analysis

Analyses were conducted using GraphPrism8 software, and the statistical tests appropriate to each analysis are indicated in Figure legends. To test for normal or lognormal distribution of sample values we used the Anderson-Darling test, outliers were identified using ROUT method. Box-and-whiskers graphs show median values with 5-95 percentile whiskers; other data are expressed as mean ± standard deviation, unless specifically stated.

## RESULTS

To distinguish only the skin-associated body wide depot (**SAF**), we used chemical shift MRI of lower limb, with fat/water separation (**Fig. 1B**). Fat-only images allow quantitative evaluation of fat content and SAF thickness, compared to the more standard fat/water MRI images (data not shown). This methodology is focused specifically on the fat content of skin, rather than the thickness of skin, which can be measured by skin fold calipers or ultrasound (Perez-Chirinos Buxade *et al*., 2018; Storchle *et al*., 2018). These other techniques are not designed to determine only the thickness of fat.

In a previous pilot study we found that SAF thickness was highly variable from subject to subject, and surprisingly, not related to body mass index (or other indices of obesity such as waist/height) (Kasza *et al*., 2016). Here, we assessed MRI images of volunteers recorded as part of a large public health study (Study of Health in Pomerania, SHIP3), including 286 women and 236 men. We found that women had thicker SAF than men (10.05±2.45 compared to 6.27±1.89 mm; **Fig.1C**). Indeed, the SAF thickness for the top decile of women was 15.0 mm, versus 6.5 mm in the bottom decile; this translates to a total predicted weight of **24.3 kg** for women with the thickest SAF layer, versus **10.5 kg** for women with the thinnest SAF, approximated using the surface area (1.8m^2^) of the average woman (Kasza *et al*., 2016). For men, this range was 9.6 mm to 3.0 mm thick (15.5-4.9 kg).

We assessed whether the thickness of SAF depended upon general adiposity, dividing the cohorts into three, lean (BMI <25), overweight (BMI 25-30) and obese (BMI >30; **Fig. 1C**). SAF did not thicken significantly in obese or over-weight men. For women, there was a significant, if minor, increase in SAF thickness for lean versus overweight women (9.00± 1.73 compared to 10.41±2.49 mm); which did not increase further in the obese cohort. These conclusions were re-stated by correlation analysis (**Fig. 1D**), showing no relationship of BMI (or Waist/hip ratio; data not shown) with skin SAF for males, and a weak relationship for females (r=0.300, p<0.0001).

We tested the distributions of values for SAF thickness for men and women, and found that for men, the values were normally distributed; for women, the values were not, best fitting a lognormal distribution, and tailing towards thicker SAF (**Fig. 1C**, S1B,C). This implies an independent factor that promotes the accumulation of SAF in a minority of women. Perhaps surprisingly, we noted no significant correlation of SAF thickness with age, for either men or women (55 men and 57 women over 70; **Fig. 1E**).

To assess similarities and differences between mouse and human skin-associated adipose depots, we first describe their morphology and anatomy. Mouse dermal white adipose tissue (dWAT) is clearly demarcated by a muscle layer (*panniculus carnosus*), making it simple to distinguish from the subcutaneous depots of the mammary glands (females) or mammary fat pads (males) (Alexander *et al*., 2015). The most commonly studied scWAT depot in mouse is the inguinal fat pad, or iWAT; this is immediately subjacent to the dWAT layer (proximity is illustrated in **Fig. 8**).

Overall, the thickness of dWAT varies from almost zero (young C57BL/6J males or rats) to 400 μm, typical of anagen stage dorsal skin for BALB/cJ mice, or obese mice (**Fig. 2A, B, D, E**). The average thickness for non-anagen stage dorsal skin from chow-fed adult BALB/cJ females housed in room temperature housing is 350 μm (**Fig.2D**).

**Fig. 2.**
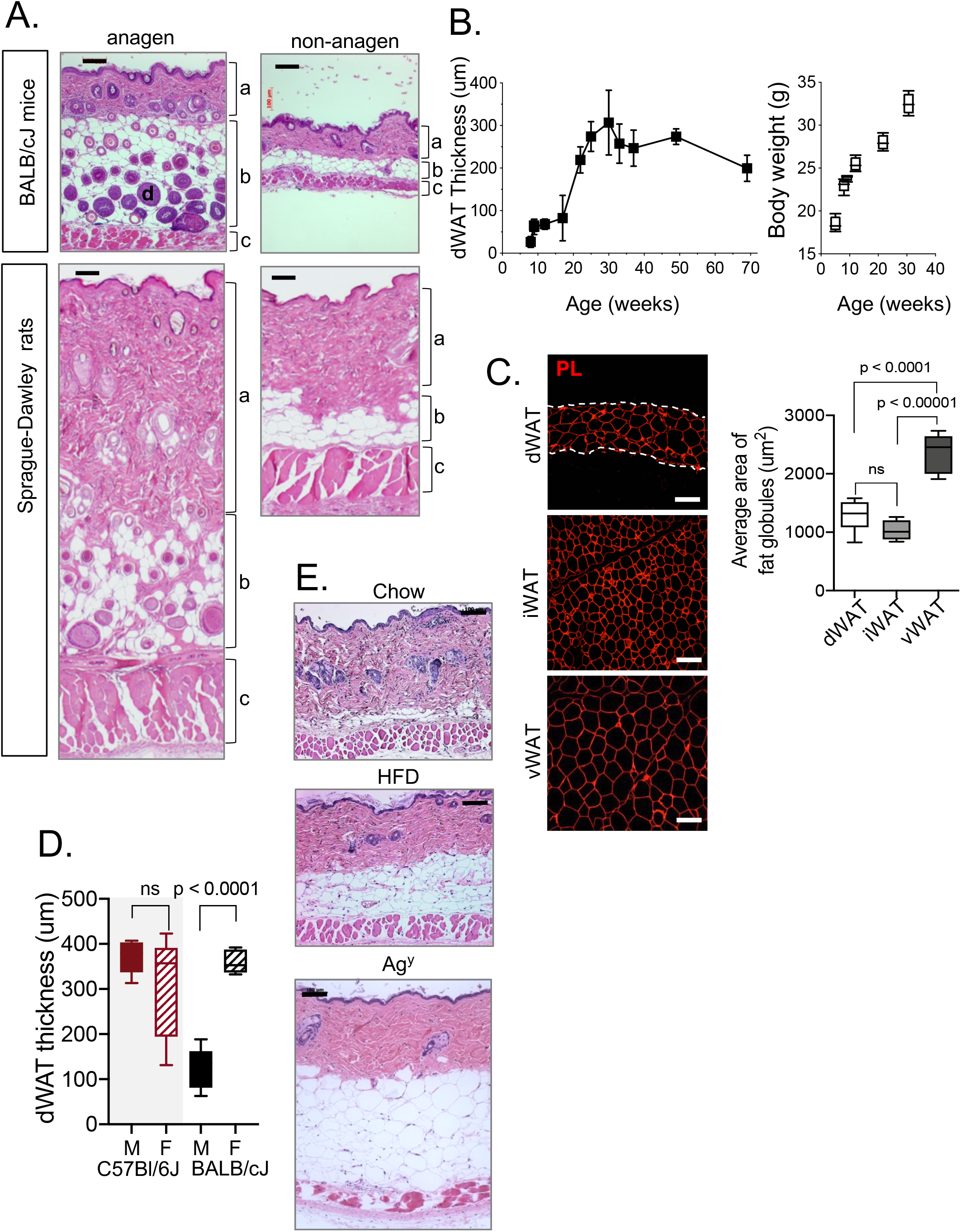
Determinants of rodent dWAT thickness; age, sex, hair cycle, and obesity. **A.** *Hair cycle associated expansion*. Representative H&E stained sections of skin of rats and mice in non-anagen and anagen (follicular development) phases. **a**, dermis/epidermis; **b**, dWAT; **c**, *panniculus carnosus*; **d**, hair follicle. Scale bars=100 μm. **B**. *Maturation dependent accumulation*. Assay of dWAT thickness during the growth and maturation of male C57BL/6J mice; n=65. **C**. *Dermal adipocytes match inguinal adipocyte size*. Comparison of the sizes of fat globules in dermal (dWAT), inguinal (subcutaneous; iWAT) and visceral (vWAT) adipocytes from lean mice, as measured by immunofluorescent visualization using anti-perilipin (PL); n=6. Scale bars=100 μm. **D**. *Thickness of dWAT is highly sex- and strain-dependent*. dWAT thickness was measured from H&E-stained sections of skins from C57Bl/J or BALB/cJ male and female mice >30 weeks old (n=6). Data of panels B and D were analyzed by unpaired two tailed *t* test; groups compared for panel C were analyzed by one-way ANOVA followed by Tukey’s multiple comparisons test. **E**. *Number and size of dermal adipocytes increases in obese mice*. Representative H&E sections of dorsal skins from C57BL/6J male mice (7-11 weeks; n=6), fed chow or high fat diet for 5 weeks, or from a genetically obese model (1 year old C57BL/6J-Ag^y^; n=3). Average diameters are shown in Table 1. Scale bars=100 μm.

We noticed that dWAT accumulates steeply in C57BL/6J males between the ages of 20 and 30 weeks; this occurs prior to, and during, a phase of rapid continuous body weight gain (**Fig. 2B**).

Using an immunohistochemical stain to outline the fat globule of adipocytes (anti-perilipin), we found that mouse dermal adipocytes contain approximately the same amount of fat as inguinal/subcutaneous WAT adipocytes but show only half the cross-sectional area compared to visceral adipocytes (**Fig. 2C**). Assuming the fat globule is approximately spherical, this translates to approximately 4-fold higher lipid load in each visceral adipocyte compared to the adipocytes in iWAT and dWAT depots.

We compared the thickness of dWAT in males and females from two strains, C57BL/6J and BALB/cJ mice, and found that BALB/cJ mice showed a dramatic sex dimorphism, where dWAT was almost 4x thicker in females than males (**Fig. 2D**). This was not observed for mature male and female C57BL/6J mice, though mature females showed high variability compared to males.

Bearing in mind the lack of correlation between human SAF thickness and obesity, we tested whether obesity and dWAT thickness were correlated in mice. We found that dWAT thickness expanded by 4-fold after 2 weeks of high fat feeding in for C57BL/6J males, in parallel with vWAT depots (**Fig. 2E** and Kasza et al 2016 (Kasza *et al*., 2016)). Likewise, genetically obese mice (A^y^/a) showed thick dWAT and increased dermal adipocyte size; diameters of dermal adipocytes (obese and lean) are summarized in **Table 1**.

**TABLE 1.**
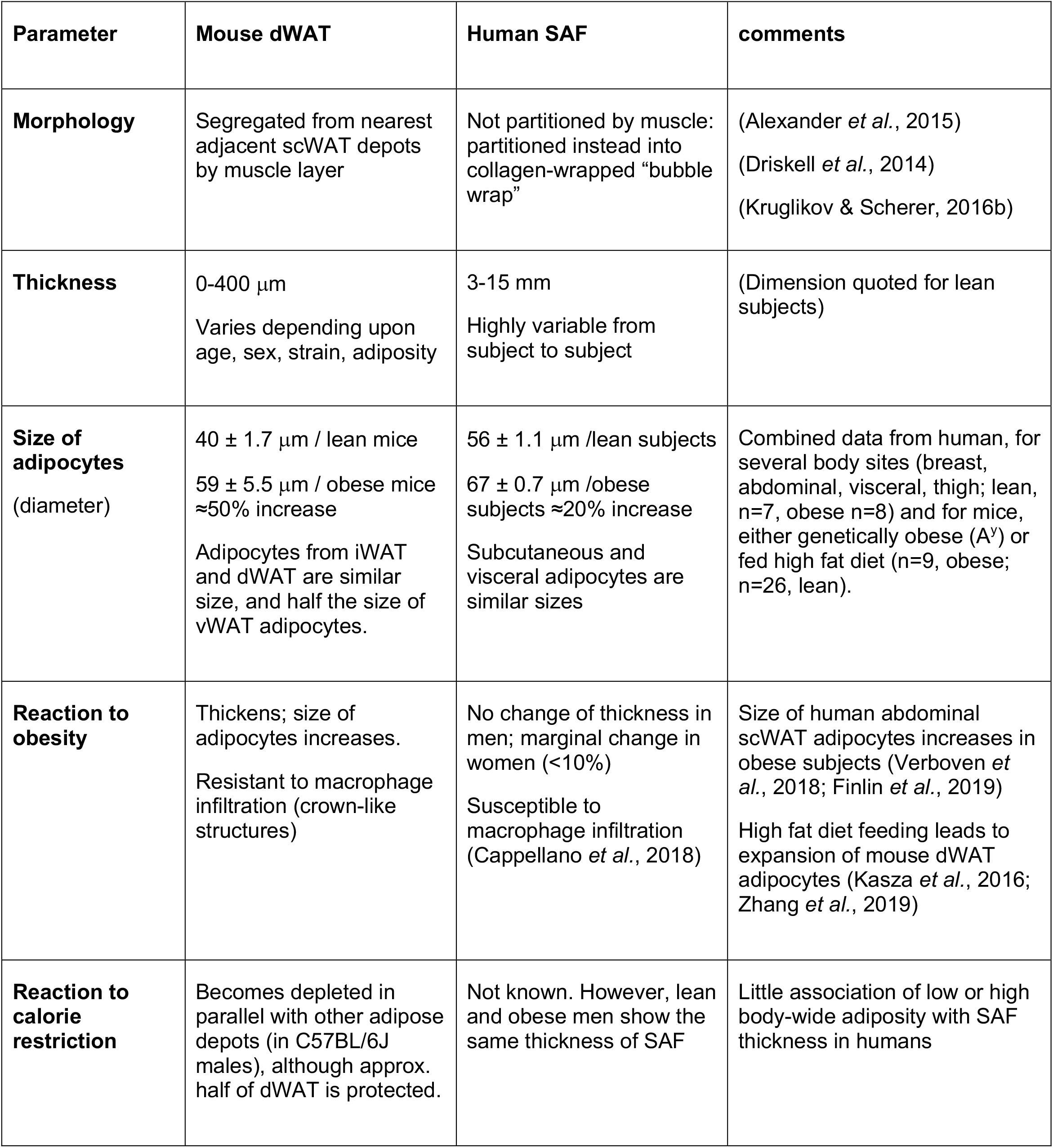
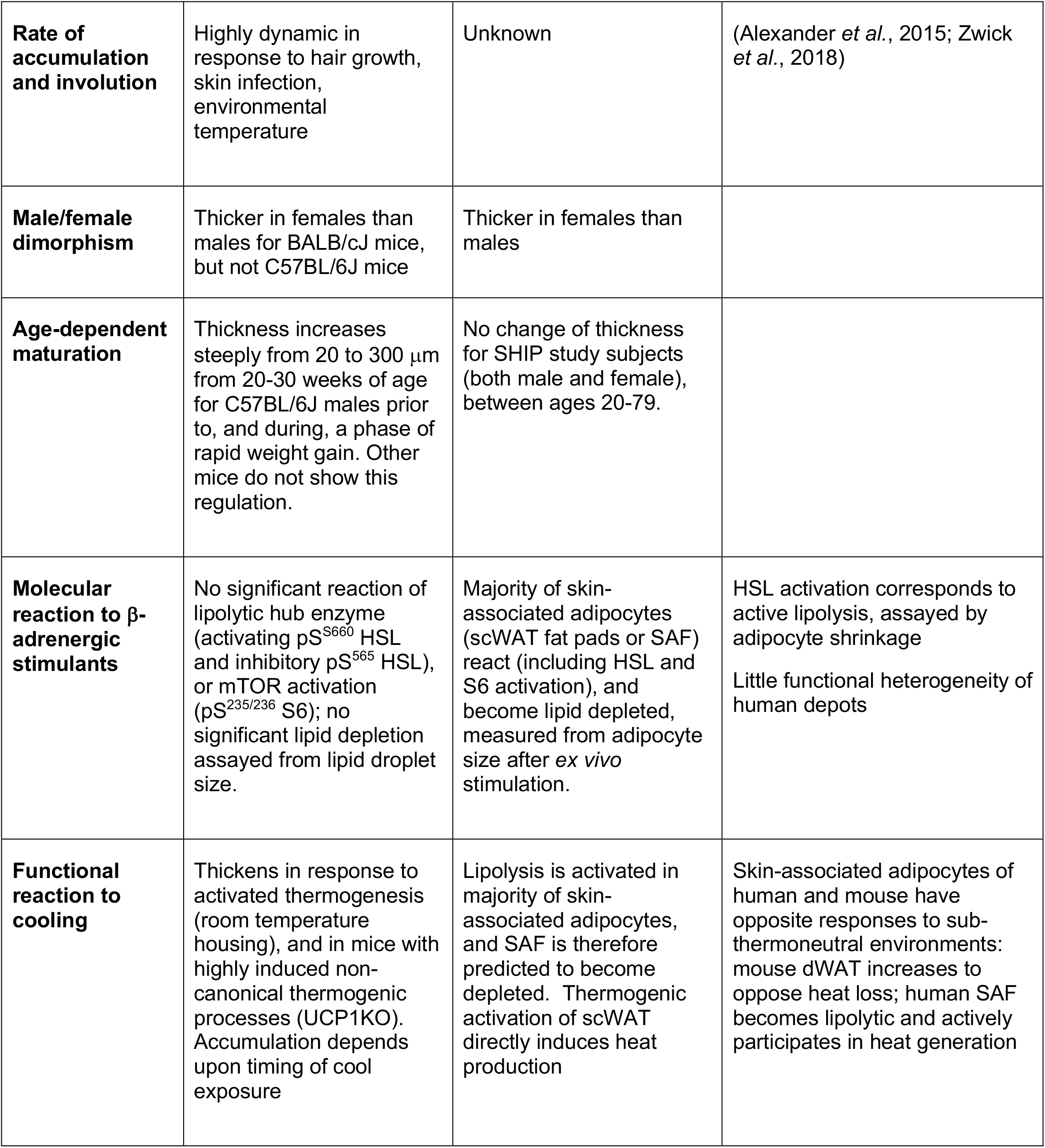
Summary comparison of key features of skin-associated fat depots for mouse and human.

Inflammation of obese adipose depots is linked to metabolic deterioration, and adipose depots from obese mice are differentially sensitive to invasion by inflammatory macrophages, assayed by the accumulation of crown-like structures (CLS) (Grove *et al*., 2010). We investigated whether CLS appeared in dWAT depots in skins from mice fed with high-fat diets (compared to either iWAT or gWAT) and found that this depot was relatively resistant to this type of inflammation. Some crown-like structures appeared in mice genetically altered to have life-long obesity (32 week-old A^y^ mice; **Fig. S2A**).

*Vice versa*, calorie restricted mice showed depleted dWAT. We used a moderate, 25% calorie restriction protocol to assess impact on both C57BL/6J and BALB/cJ mice; this protocol induced a significant loss of body weight within 2 weeks. After 4 weeks of calorie restriction, C57BL/6J mice lost 18% body weight, and BALB/cJ mice lost 22% body weight (**Fig. 3A**). DWAT mirrored the loss of body weight; both strains lost >100 μm thickness of dWAT (**Fig. 3B**), so the leanest calorie restricted BALB/cJ mice showed negligible dWAT levels after 4 weeks (**Fig. S2B**).

**Fig. 3.**
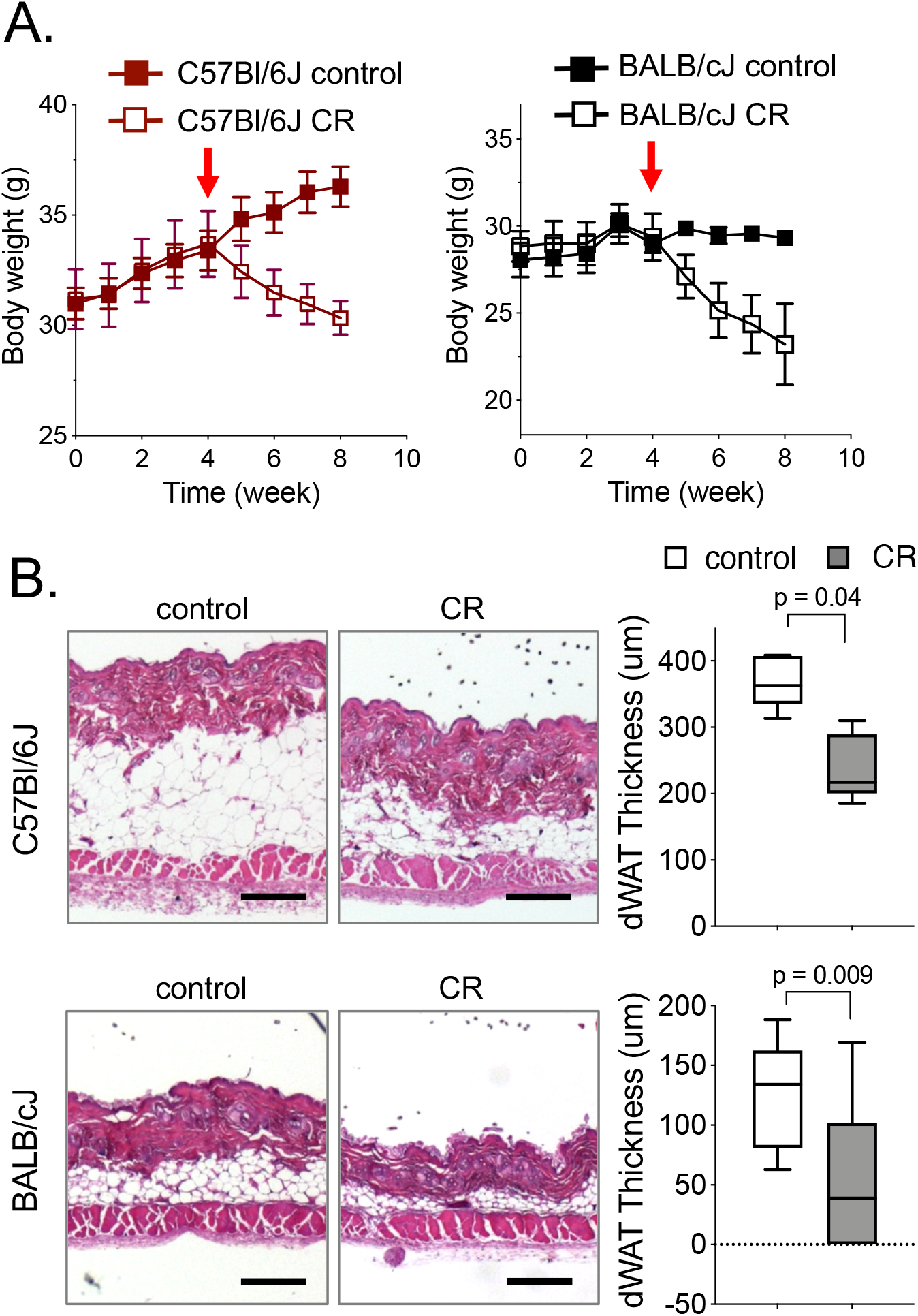
Calorie restriction depletes dWAT. **A.** Body weight changes for C57BL/6J or BALB/cJ male mice (28 to 30-week-old) after 25% calorie restriction for 4 weeks (red arrow indicates start of dietary regime); n=6. **B**. dWAT thickness was assessed from H&E sections of dorsal skin. Scale bars=300 μm. Data were analyzed by unpaired two tailed *t* tests.

In contrast, human subcutaneous adipose tissues comprise capsules of yellow fat attached to the dermis/epidermis; capsule size ranges from 5-15 mm, where the lipid-filled adipocytes are encapsulated by thick seams of collagen (**Fig. 4A**). Several publications have reviewed the topic of potential heterogeneity of human subcutaneous adipose tissue (Kelley *et al*., 2000; Smith *et al*., 2001; Sbarbati *et al*., 2010; Kruglikov & Scherer, 2016a; Zwick *et al*., 2018); in particular, deep layers of skin-associated fat may have different properties compared to the more superficial layers (Enevoldsen *et al*., 2001; Cappellano *et al*., 2018). We found little difference in the size of adipocytes from superficial (within 5 mm of dermis), mid (5-15 mm) and deeper layers (more than 15 mm from dermis; n=7) for samples from lean individuals (**Fig. 4A, B**); neither did we observe significant differences in adipocyte sizes from different body locations (for example breast versus lower body (thigh) versus upper body (abdominal); n=7, 4, 5 respectively) (**Fig. 4B, C**). The adipocytes from visceral depots and breast and abdominal subcutaneous sites all underwent hypertrophy (**Fig. 4C**). In summary, human subcutaneous adipocytes of lean subjects were larger on average (56 μm diameter) than mouse dermal adipocytes (40 μm diameter), and adipocytes from both human and mouse undergo hypertrophy during obesogenesis (67 and 59 μm respectively; **Fig. 4D; Table 1**).

**Fig. 4.**
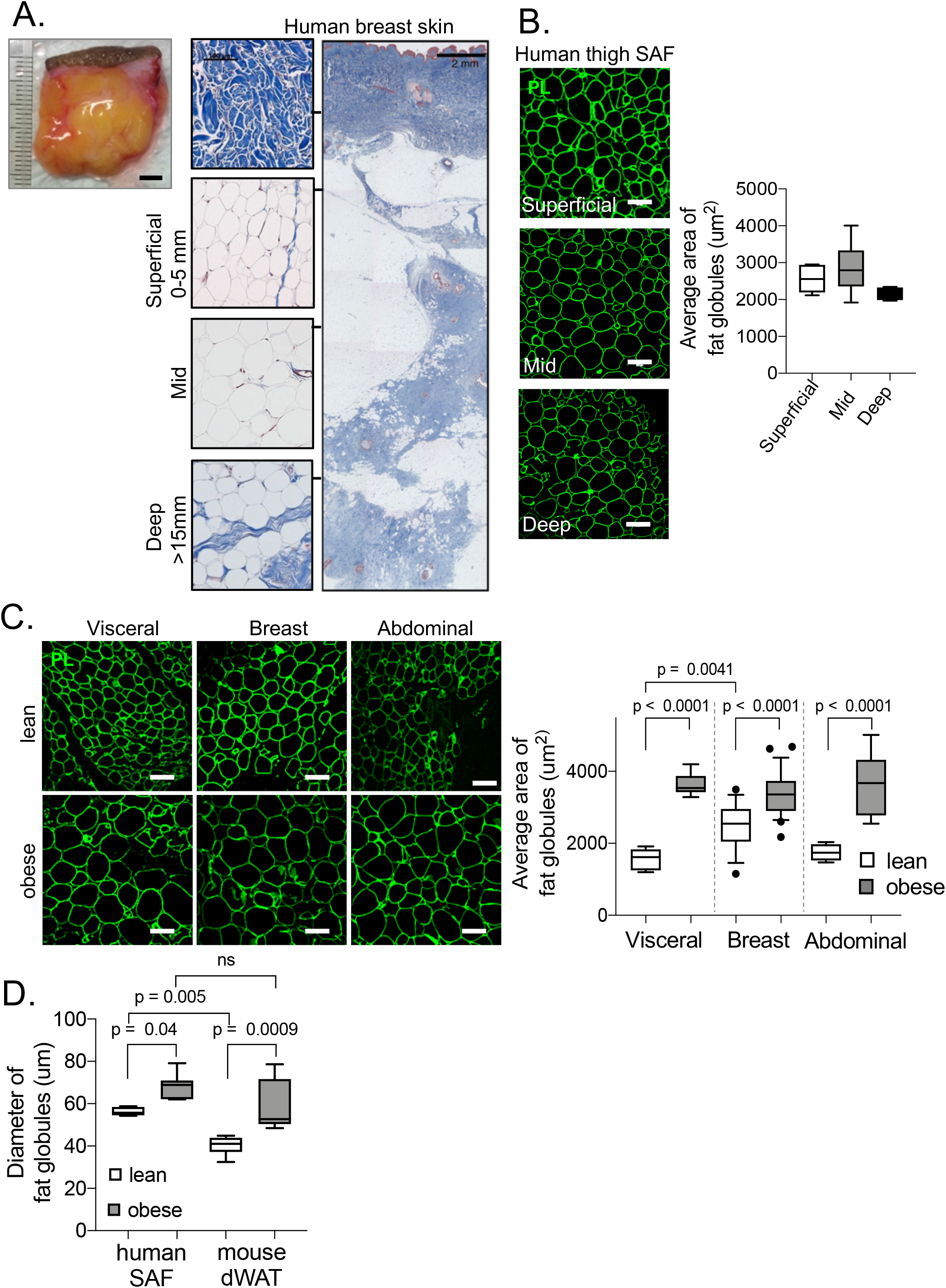
Like mouse dermal adipocytes, human skin-associated adipocytes hypertrophy in obese subjects. **A.** *Gross anatomy and histology of human skin-associated fat*. Gross appearance and trichrome-stained sections of SAF from breast of a lean subject, with detail insets shown for dermis, superficial, mid and deep layers of adipocytes. Scale bars=5 mm on gross view, 2 mm on Trichrome-stained low power view, and 100 μm on high power detail insets. **B**. *Size of adipocytes is unaffected by distance from skin*. Human thigh tissue sections were stained with perilipin antibody (n=3); representative images are shown, with quantitation of the size of adipocytes from superficial, mid and deep layers. Scale bars=100 μm. **C**. *Comparison of SAF from breast and abdomen with visceral WAT*. Representative images of PLIN-stained fat globules from lean and obese subjects (left) were quantified (right); scale bars=100 μm. **D**. *Adipocytes of mouse dWAT and human SAF respond similarly to obesogenesis*. The relative size of adipocytes in human SAF and mouse dWAT are shown for lean and obese subjects (human lean, n=5; human obese, n=7; mouse lean, n=7; mouse obese, n=5). Data were analyzed using a one-way ANOVA followed by Tukey’s multiple comparisons test.

Human and mouse subcutaneous adipose depots are typically ascribed a supportive role in the thermogenic response; they are activated by β-adrenergic agents to become lipolytic, to provide fatty acid fuels for heat production by BAT, or to participate in the beiging response (Wu *et al*., 2012; Mottillo *et al*., 2014; Chondronikola & Sidossis, 2019). However, the analysis of the transcriptome of mouse dermal adipocytes performed by Scherer and colleagues suggested that this population was unlike adipocytes in brown or beiging depots, resembling vWAT instead (Zhang *et al*., 2019); they showed that dWAT showed no lipid depletion upon treatment with a β-adrenergic agent (scored as a reduction from mono-to multi-locularity of the lipid body).

To further evaluate the molecular response of dermal white adipocytes to cold stress, we evaluated the regulatory modification of hormone-sensitive lipase (HSL) in response to the β-adrenergic agonist CL *in vivo* using an immunohistochemical assay (**Fig. 5**). HSL is part of a complex that integrates input stimuli from kinases to coordinate lipolysis brown adipose tissue (Fruhbeck *et al*., 2014; Ogasawara *et al*., 2015). We confirmed the accuracy of HSL (and FASN) stains using mouse adipocyte cultures treated with the pan β-adrenergic agonist, isoproterenol (**Fig. S3**). Tissue samples of BAT and iWAT showed a gain of signal for the PKA-dependent activating phosphorylation of S^660^ (**pHSL stim**), whereas vWAT showed a gain of signal for the AMPK-dependent inhibitory phosphorylation of S^565^ (**pHSL inh**; **Fig. 5A, B**). Positive staining for fatty acid synthase, FASN, and fatty acid binding protein-4, FABP4, was suppressed in iWAT and BAT, and unchanged/negative for vWAT and dWAT (**Fig. S4**). This confirms that dWAT was non-responsive to β-adrenergic-induced heat production, measured by the same criteria used to assess the recruitment of brown and beige depots.

**Fig. 5.**
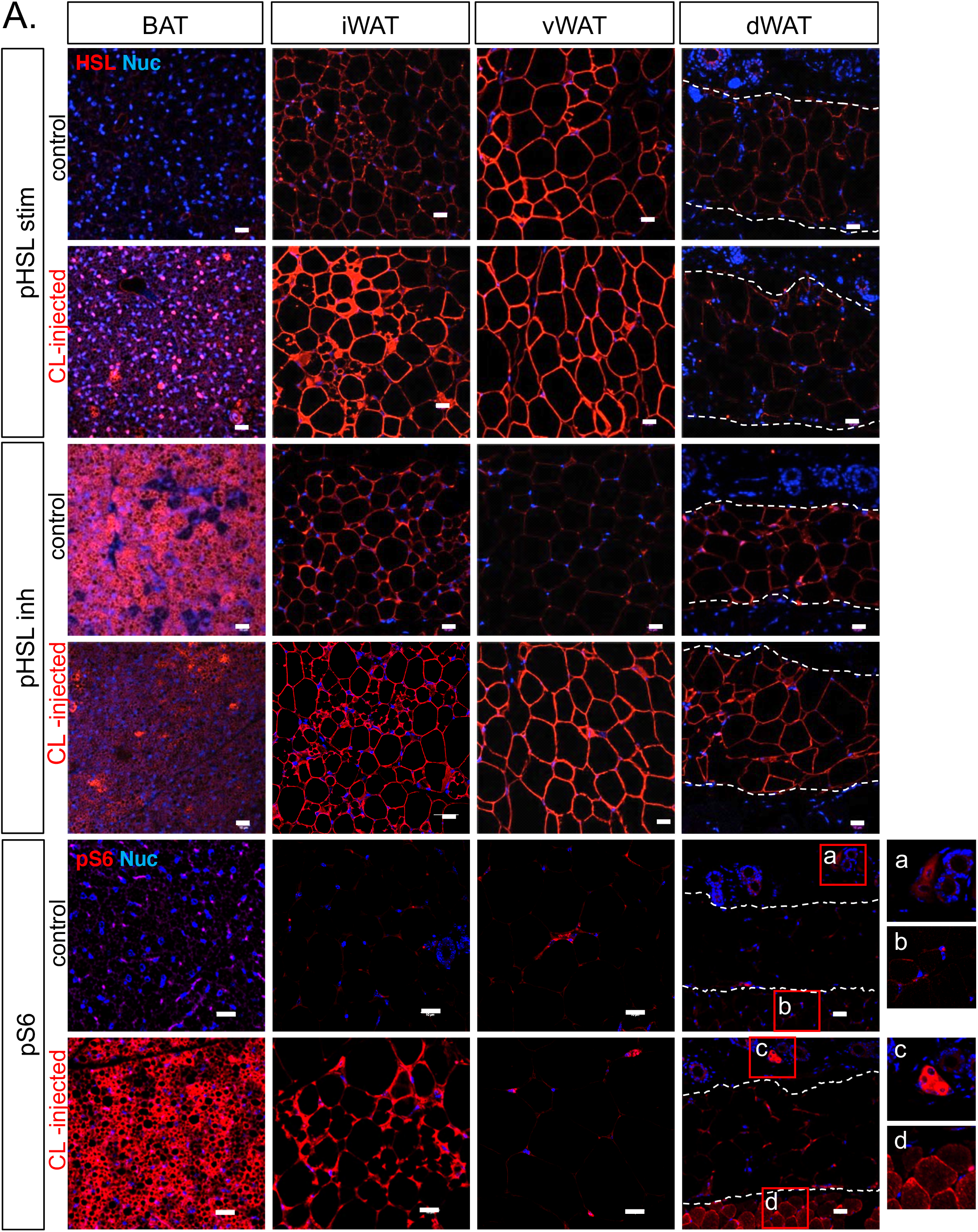
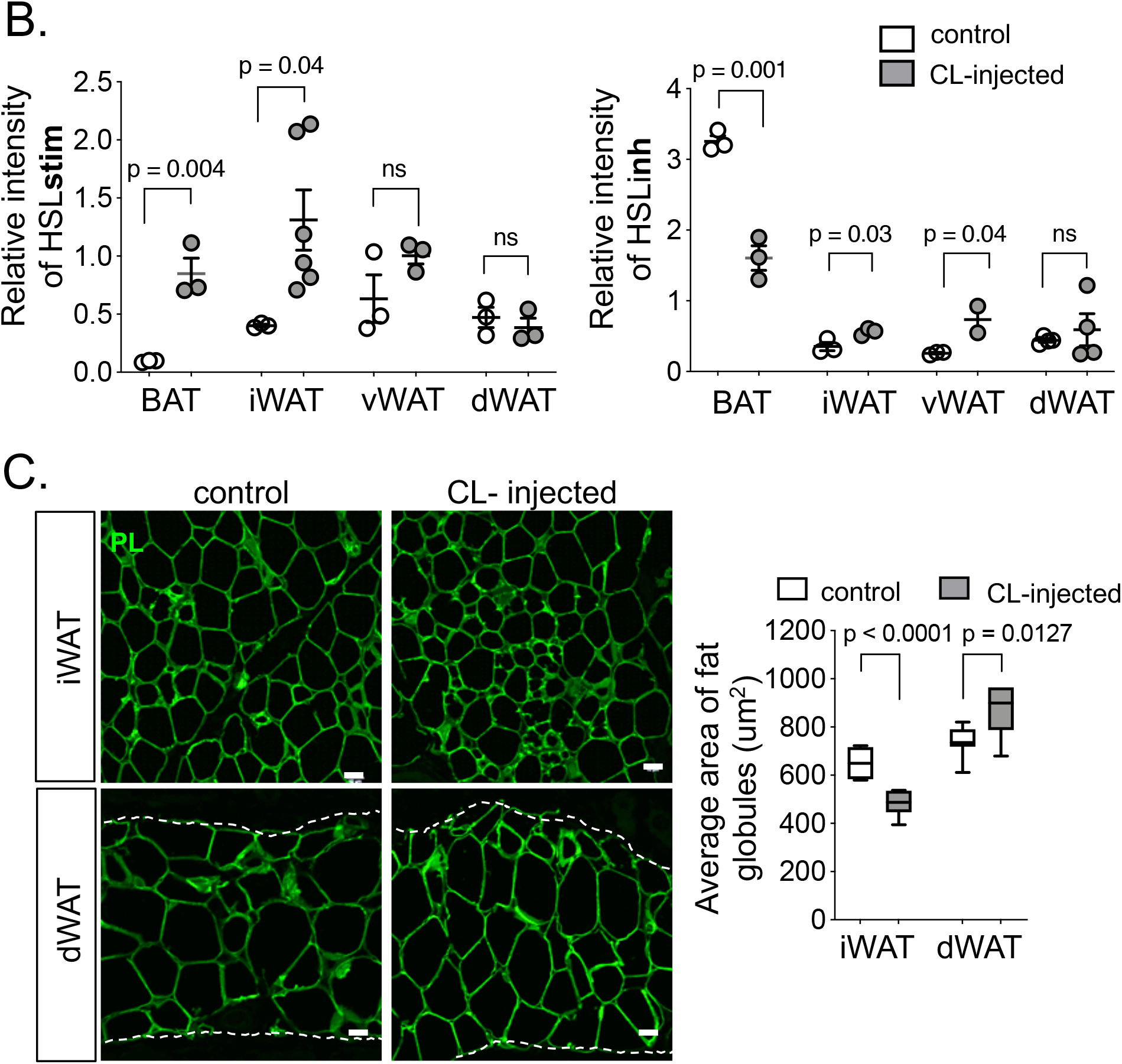
Mouse dWAT is entirely insensitive to β-adrenergic lipolysis. **A.** *HSL and mTOR are not activated in dWAT after treatment with a β-adrenergic agonist*. Fat depots (brown adipose tissue, BAT; iWAT, vWAT and dWAT) were harvested from mice, euthanized 60 minutes after administration of CL (control n=5 and treated n=4). Immunofluorescent assay of regulatory modifications of hormone-sensitive lipase (HSL), either pS^660^(pHSL stim) or pS^565^ (pHSL inh), or mTOR-activated S6 (pS^235/236^) were quantified in **(B)** as described in Methods. Insets shown for pS6-stained dWAT show sebaceous glands (a, unstimulated; c, stimulated) and *panniculus carnosus* muscle layer (b, unstimulated; d, stimulated); scale bars=20 μm. **C**. *Adipose globules of dWAT are not depleted after CL administration*. The size of lipid droplets (outlined by anti-perilipin staining) was assayed in dWAT and iWAT, 60 minutes after CL injection; scale bars=20 μm. Data were analyzed by unpaired two tailed *t* tests.

To corroborate this result, we tested another β-adrenergic/PKA-induced reaction; thus, activation of mTORC1 is known to be essential for β-adrenergic induced lipolysis in adipocytes (Liu *et al*., 2016). S6 and S6-kinase are activated and phosphorylated downstream of the mTORC1 kinase and are markers of mTORC1 activation status. Signal for phospho-S^240/244^ S6 was induced in BAT and iWAT in CL-administered mice but showed no reaction in dWAT (**Fig. 5A**), despite the local activation of muscle and sebocytes by β-adrenergic stimulation. To confirm that these immunohistochemical assays reflect active lipolysis, we measured the size of adipocyte lipid stores in mice that were activated to produce heat, either by transfer to 4^0^C environment or 60 minutes after CL administration. We found lipid stores significantly depleted in iWAT, but not in dWAT (**Fig. 5C, S5**). In summary, when BAT and iWAT become lipolytic in response to β-adrenergic stimulation, dWAT shows little molecular response, like vWAT and bone marrow adipocytes (Scheller *et al*., 2019).

We hypothesized that in mice chronically stimulated to produce more heat, dWAT would instead react to mitigate heat loss. Mice housed at room temperature (19^0^C) compared to thermoneutral housing (31^0^C) show thicker dWAT (1.6x; **Fig.6A**). To test this hypothesis using a genetic model, we evaluated *UCP1-/-* mice. These mice have a lesion in the most efficient heat production mechanism (affecting the uncoupling of BAT mitochondria), and deploy instead beiging and other “non-canonical” mechanisms to compensate (Keipert *et al*., 2020). As with environmentally cool-stressed mice, genetically cool-stressed mice also show thicker dWAT (**Fig. 6B**). This supports the claim that the accumulation of fat in dWAT is an adaptive response to environmental cooling.

**Fig. 6.**
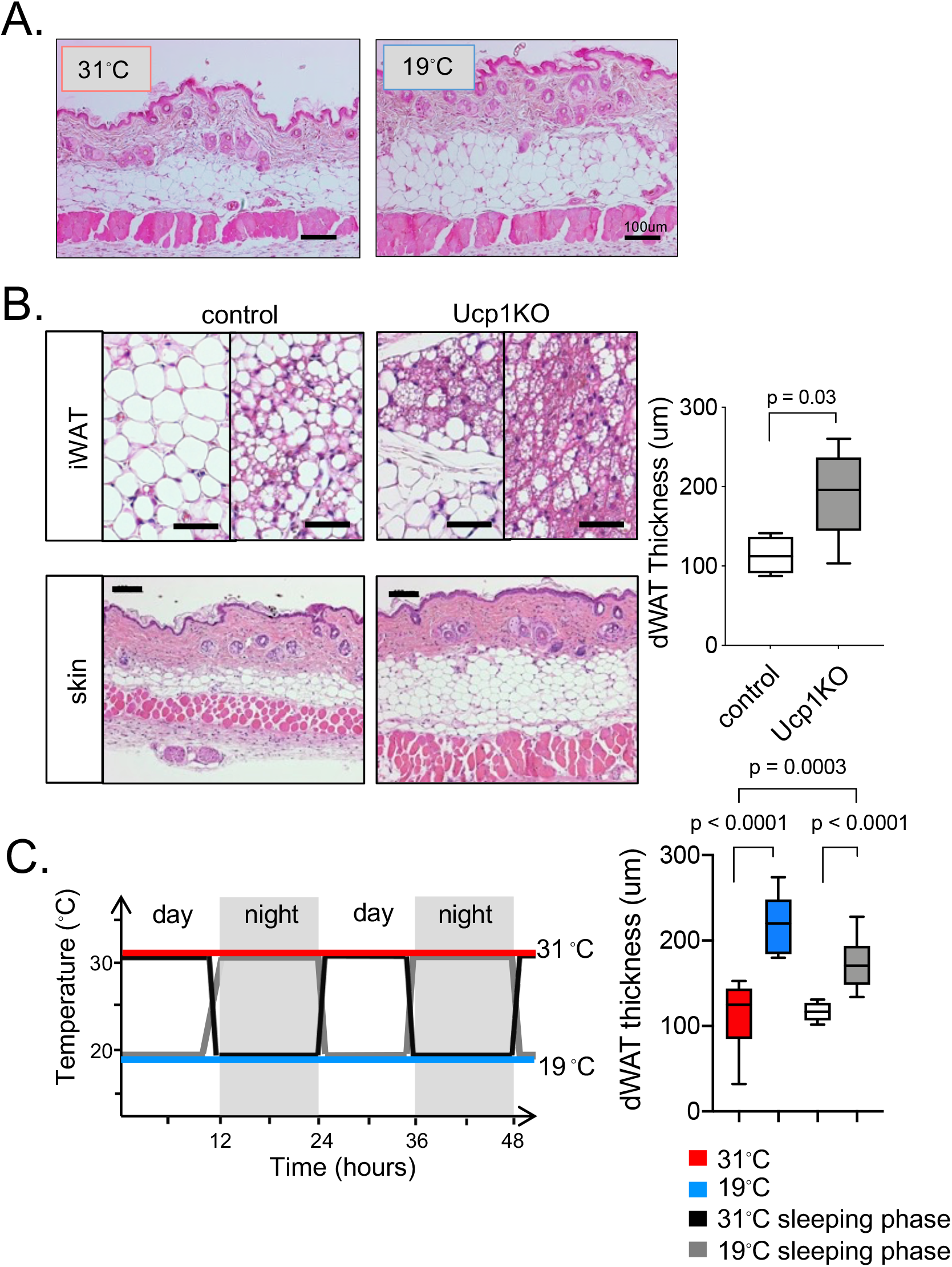
Mouse dWAT accumulates when mice are challenged to produce heat. **A.** *dWAT thickens in response to sub-thermoneutral environmental housing*. dWAT thickness was assayed for mice housed at thermoneutrality (31^0^C) or room temperature (19^0^C) for 3 weeks; scale bars=100 μm. **B**. *dWAT thickens in UCP1-/- mice producing heat by non-canonical means*. H&E stained sections are shown for iWAT and dWAT from *UCP1-/-* or control C57BL/6J mice (10 to 19-week-old); quantified at right hand side. Scale bars=50 μm (iWAT) or 100 μm (skin); n=≥4. **C**. *Circadian cues promote dWAT accumulation*. Mice were housed individually with alternating circadian cycles of thermoneutral and room temperature housing for 3 weeks. The experimental paradigm is shown on the left, continuous warm (31^0^C, red line), continuous cool (19^0^C, blue line), or alternating daily cycles of warm sleeping /cool waking phases (black) or cool sleeping/warm waking phases (grey). The thickness of dWAT is shown on the right (n=5). Data were analyzed by unpaired two tailed *t* tests.

What triggers this adaptive response? In particular, we noticed that mammals typically experience variation of environmental temperature over the circadian daily cycle. To begin to isolate environmental cues, we divided the housing cycle into a warm phase (thermoneutral, 31^0^C/88^0^F) and a cool phase (cool room temperature, 19^0^C/66^0^F). Although the length of time in each phase was constant for mice in cycling environments, only the mice housed cool during their sleep cycle developed thicker dWAT (**Fig. 6C**); the same cue given to awake mice did not induce dWAT thickening. This suggests that there is a component of circadian control of dWAT lipogenesis.

Turning to the molecular regulation of skin-associated fat in human, we administered β-adrenergic agonists either *in vivo* or *ex vivo*, and assayed HSL activation and lipid store mobilization. Human abdominal, breast and thigh skin-associated fat administered the pan-β adrenergic agent isoproterenol *ex vivo* showed uniform activation of HSL and lipid store depletion (**Fig. 7A, B**). Likewise, abdominal adipose tissues from obese subjects administered the β3-adrenergic agent mirabegron daily, also showed activated HSL, and higher UCP1 staining (**Fig. S6**)(Finlin *et al*., 2018). Skin-associated fat in calf, thus the same body site imaged for the data of Fig. 1, also showed HSL activation after exposure to isoproterenol (**Fig. S7**). This β-adrenergic response was therefore a consistent feature of human skin-associated adipose tissues, regardless of body site or distance from skin (superficial or deep).

**Fig. 7.**
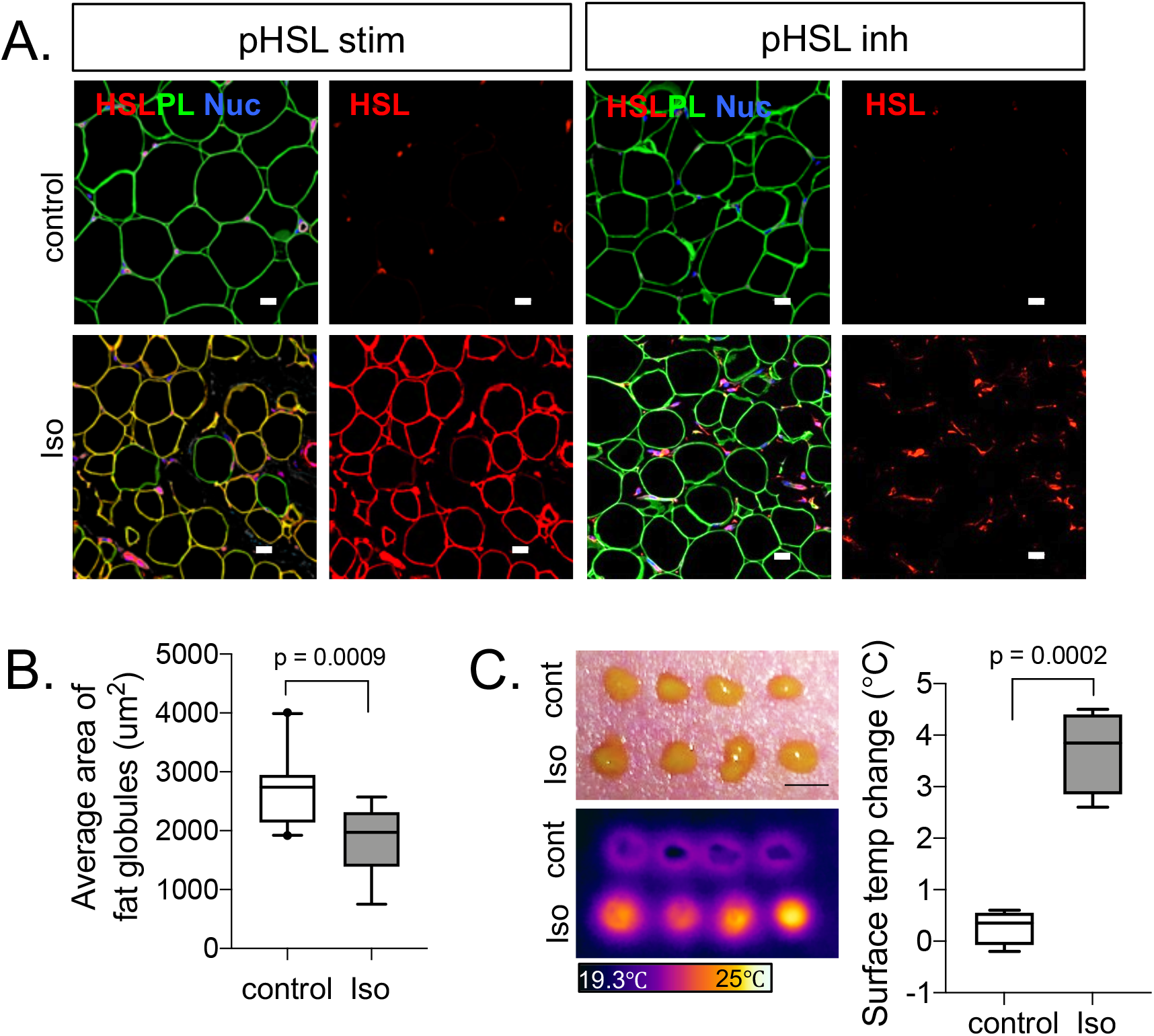
Evaluation of β-adrenergic lipolytic response for human skin-associated fat depots. **A.** *Human skin associated adipocytes all respond to β-adrenergic agonists*. Immunofluorescent stains of pHSL-stim and pHSL-inh in sections of skin-associated fat from thigh, exposed to isoproterenol (Iso) *ex vivo*, counterstained with perilipin (PL) and DAPI (nuc; nuclei) (n=3). Scale bars=20 μm. **B**. *Human skin associated adipocytes are depleted by thermogenesis*. The average size of lipid droplets was quantified for the adipocytes visualized in A. **C**. *Human skin associated fat makes heat upon treatment with β-adrenergic agonist*. Infrared thermographic images of freshly collected fat capsules, pre-treated or not with isoproterenol for 40 mins, were quantified (right hand side). Scale bar=5mm. Data were analyzed using unpaired 2-tailed t tests.

**Fig. 8.**
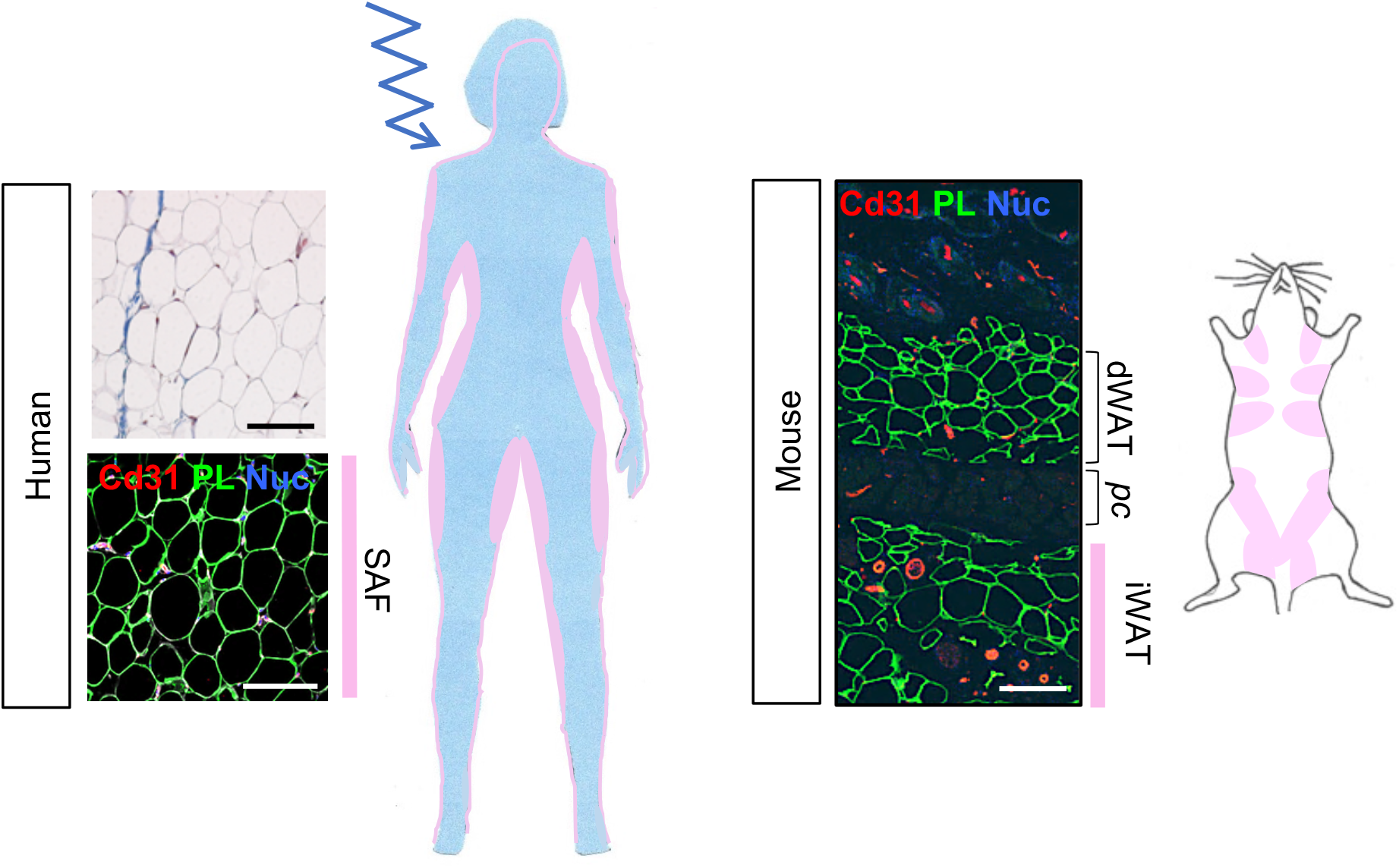
Summary of inferred functionality of skin-associated adipose depots for human and mouse. In human (left hand side), the skin-associated fat depot is responsive to β-adrenergic demand and contributes heat towards body temperature homeostasis. For mouse (right hand side), there is a bilaminar adipose depot, the superficial dWAT layer is non-responsive to β-adrenergic demand (increasing insulative properties), whereas the subjacent iWAT depots are responsive. Immunofluorescent stained sections (both human and mouse) are stained for endothelial cells (CD31), perilipin (PL) and DAPI (nuc; nuclei); scale bars=100 μm.

Typically, lipolysis of subcutaneous fat is considered to have a supporting role for the delivery of fatty acids to dedicated, heat-generating organs such as brown adipose tissue or muscle. However, emerging literature suggests that a number of UCP1-independent biochemical pathways can be harnessed for the purpose of regulated heat production (Chouchani *et al*., 2019), where the substrates used to fuel oxidation are not yet clearly defined. Therefore, to directly evaluate whether β-adrenergic activation can induce heat production in skin-associated fat, we used infrared thermography (FLIR) to assay fat explants *ex vivo* and found a significant rise in temperature in response to isoproterenol treatment (**Fig. 7C**). We conclude that human skin-associated adipose tissue is an active participant in heat production, and not just a passive insulator.

## DISCUSSION

We compared the most superficial layers of adipose associated with skins of human and mouse and showed that they respond differently to the activation of thermogenesis; thus the dermal adipocytes associated with mouse skin are protected from lipolysis, where the skin-associated fat of human skin becomes lipolytic and produces heat autonomously (**Table 1**). Thus, the human skin-associated fat layer comprises a dynamically warming blanket, not only able to resist heat loss, but actively contributing to the total heat budget (see scheme in **Fig. 8**). Our data show that human skin-associated adipocytes are remarkably homogeneous in their ability to recognize and respond to β-adrenergic effectors, whether from obese or lean subjects, and irrespective of body site (upper versus lower body, or limb versus abdominal depots). A speculative review has suggested that there could be a dWAT equivalent around human hair follicles (Kruglikov & Scherer, 2016a), but we could find no β-adrenergic-resistant adipocyte population in the array of human samples we evaluated. Note that we use the umbrella terminology of “skin-associated fat” for human, aware that there is an extensive literature documenting site-specific regulation and properties of subcutaneous adipose tissues.

This study is the first to quantify this specific human fat depot, though many MRI-based studies have reported segregated fat volumes that discriminate between, for example, visceral and sub-cutaneous abdominal fat (Wajchenberg, 2000; Smith *et al*., 2001). Specific “fat-only” MRI data can be extracted from the data obtained for routine diagnostics, though those scans are typically processed to show a combination of fat and water signals. Other techniques aimed at assessing body composition use ultrasound or calipers to measure skin-fold thickness (Perez-Chirinos Buxade *et al*., 2018; Storchle *et al*., 2018), however, although these techniques may show broadly comparable results, they do not measure fat specifically.

Other imaging strategies used for the study of thermogenesis rely on the detection of glucose uptake by ^18^F-fluorodeoxyglucose (FDG)-PET imaging of activated fat depots in cool-exposed individuals. The SAF depot has not yet been noted by these studies (Chen *et al*., 2016). This could be due to 1) the relatively lower resolution of this technique and the disseminated nature of the depot, or 2) low glucose uptake by activated SAF. This is possible, since we do not yet understand the substrate requirements of heat production in non-BAT thermogenic tissues, but they probably include creatine and amino acids (Ikeda *et al*., 2017; Mills *et al*., 2018; Kazak *et al*., 2019).

Women show thicker SAF than men, by 60% on average; by calculation this depot is 10.5-24.3 kgs, making it the largest individual adipose depot in the lean woman’s body. The number of women with thick SAF is higher than that predicted from a normal distribution, and we speculate that the health and energy expenditure of this population may be significantly affected. Likewise, the females of the BALB/cJ mouse strain show much thicker dWAT than their male counterparts; this depot is thus more prominent in females of both species.

The dWAT depot of mice correlates strongly with relative adiposity elsewhere, even from mouse to mouse in a given population. Mouse dWAT increases in obese mice, whether induced by high fat feeding, by age-induced obesity, or in mice made obese genetically, including *Agouti*^*y*^, MitoNEET and *ob/ob* ((Kasza *et al*., 2016; Kasza *et al*., 2019; Zhang *et al*., 2019) and this study). In adult mice, the majority of dWAT thickening is enabled by adipocyte hypertrophy rather than hyperplasia (Zhang *et al*., 2019).

In obese mice, we found this depot to be relatively resistant to the development of “crown-like structures”; these reflect the infiltration of inflammatory macrophages into depots containing distended adipocytes, typically highest in male mouse vWAT (Grove *et al*., 2010). For human subjects, subcutaneous adipose depots are significantly associated with inflammatory macrophage infiltration (Cappellano *et al*., 2018).

*Vice versa*, we showed that dWAT is depleted in mice subjected to calorie restriction. Trajkovski and colleagues showed that beige depots were activated in calorie restricted mice (Fabbiano *et al*., 2016); this may support the hypothesis that dWAT thinning leads to higher demand for heat.

Given *ad libitum* calorie intake, we propose that mice deploy the combination of iWAT activation and dWAT thickening to meet heat production demands, notably when BAT activation is insufficient. Thus UCP1KO mice deploy a range of strategies for heat conservation and production (Chouchani *et al*., 2019), to ensure their body temperature is resilient to change upon cool stress. As well as thicker dWAT (shown here), UCP1 KO mice show a change of vascular tone, so mice have a sustained vasoconstrictor response after minor cool stress (Kajimura *et al*., 2015). Fgf21 was shown to be an effector of iWAT activation in UCP1KO mice (Keipert *et al*., 2020); perhaps Fgf21 is also an effector of dermal adipocyte hypertrophy. We note that the dWAT depot, alongside other fat depots, becomes depleted when mice are challenged with an extreme cold stress (transfer to 4^0^C)(Zhang *et al*., 2019), which could compound their energetic stress.

This study has shown little correlation between SAF thickness in human subjects and obesity, indeed the thickness of this layer is highly individual specific. The marginal increase in thickness between lean and overweight/obese individuals could be accounted for by the 20% increase in SAF adipocyte fat globule size (**Table 1**). We speculate that both environmental and genetic factors may control the thickness, with impact for subsequent energy budget. Specific genes are known to regulate the pattern of human fat deposition, notably for example lower body subcutaneous depots (such as gluteofemoral) and peritoneal visceral fat (Loh *et al*., 2015; Lu *et al*., 2016); there may be similar genes that regulate the thickness of SAF.

We also note that high variation in SAF volume may lead to a correspondingly high variation in the total heat generating capacity of this depot. Heat production in mammalian bodies is a zero-sum reaction, the demand for heat is fulfilled by various means, whether muscle exercise, activation of BAT, or activation of non-canonical futile cycles in beige fat, muscle or heart. This leads us to speculate that our observations could go some way to explaining the paradoxical differences observed for individual lean subjects in their capacity for BAT activation (van Marken Lichtenbelt *et al*., 2009); BAT activation for any given individual may be inversely related to the total amount of skin-associated adipose tissue. It would also explain why obese subjects show little BAT activation, since these individuals typically have a large volume of subcutaneous fat pads.

We considered the possibility that the lipolytic hub, HSL, was not present to be activated in dWAT in response to β-adrenergic activation; however, activation of this enzyme has been observed during the lipid depletion associated with dWAT involution during catagen (Rivera-Gonzalez *et al*., 2016). The lack of HSL activation in dWAT stands in contrast to human SAF, where it is known that lipolysis is dependent upon HSL (Langin *et al*.,2005). Indeed, human scWAT has been shown to become lipolytic during the cool season (Kern *et al*., 2014; Finlin *et al*., 2017; Finlin *et al*., 2018), when mast cells are implicated as the lipolytic initiator: Only temporary but repeated application of an ice pack to thigh (30 minutes for 10 days) was enough to induce systemic activation of the β-adrenergic response. We would expect sustained lipolysis to deplete the SAF layer (Fig.7B); it remains to be tested whether intermittent cool exposure increases or decreases the SAF layer or affects an individual’s capacity for non-canonical heat generation by subcutaneous fat activation.

Mouse dWAT is highly dynamic compared to other fat depots (Alexander *et al*., 2015), however, the time required for dWAT thickening in response to environmental cues is still on the order of days. We propose that dWAT thickening is a chronic adaptive response that mitigates heat loss when thermogenic demand is high. Acclimation to cold exposure reflects adaptive changes to heat production strategies; these are important, since they impact glucose homeostasis and insulin sensitivity (van der Lans *et al*., 2013; Hanssen *et al*., 2016; Yoneshiro *et al*., 2016). To narrow down the degree of exposure required to cue the thickening of dWAT in response to cooler environmental temperature, we exposed mice to room temperature for 12 hours out of each day, either during the waking period (night-time) or sleep period (daytime). Interestingly we found that the temperature of the mice during their sleep phase cued the development of thicker, cold-adapted dWAT, and we surmise that there is a component of circadian control of this process.

Note this cycling protocol was applied to singly housed mice; the typical sleeping temperature for group-housed mice is predicted to be warm, since mice build nests and huddle together to sleep. We conclude that our warm-sleep cycling protocol may be mimicked by group-housed mice. This implicates behavior as a regulator of dWAT thickness. The cycle of warm-sleep and cool-waking time was designed to mimic a typical human circadian environmental temperature exposure, with the goal of identifying whether there are dominant temperature exposures that train the thermogenic response of each individual mammal. Our data to date suggests that chronic adaptations are designed to minimize the total thermogenic load, therefore intermittent exposure to cool temperatures will likely have more benefit than continuous chronic exposures, when homeostatic mechanisms are activated.

Scherer and colleagues defined aspects of mouse dermal adipocytes that make them unique, such as microenvironment, transcriptome and regulation. This group found that these adipocytes express distinguishing gene markers such as cathelicidin (CAMP1), collagen5 and CCL4 (Zhang *et al*., 2019). We have shown that the most distinctive property of dWAT, that distinguishes it from its nearest neighbor, iWAT, is its ability to resist depletion in response to β-adrenergic activation (**Fig. 5; Table 1**).

Despite the distinct properties of human SAF and mouse dWAT, there is a functional similarity between the heat production strategies of mouse and human skin-associated depots. Thus, directly underneath the dWAT heat transfer barrier lie ten subcutaneous fat pads (mammary glands or male equivalent; **Fig. 8**) spread across the peritoneum, where the approximate surface area equals 10cm^2^. These are highly responsive to β-adrenergic activation (including heterogeneous UCP1 induction), and we conclude that together, the dWAT and iWAT combination are functionally homologous to human skin-associated fat.

## ADDITIONAL INFORMATION

### Author contributions

**CMA** Development of strategy, execution of experimental procedures, consultation with analysis, manuscript preparation; **IK** Design and execution of experimental procedures, data analysis and presentation, manuscript preparation**; JK, HZ, RB, DH** Design and execution of MRI imaging procedures, data analysis and presentation**; AG, YZ, JS, PK** Collection of human skin and fat samples; **CLEY, DN, NR, DL** Development of strategy, consultation on mouse metabolic studies, supply of samples from mice on high fat feeding protocols**; OM** Development of strategy, manuscript preparation. All authors approved the final version of this manuscript and agree to be accountable for the accuracy and integrity of the work.

### Data Availability Statement

This article is published at BioRxiv, including supplemental data confirming the accuracy of reagents used in this study: https://www.biorxiv.org/content/10.1101/2020.09.16.300533

Full datasets are described in the Statistical Summary.

## Acknowledgements

We appreciate expert technical assistance from Edgar Ocotl, the University of Wisconsin Translational Research Initiatives in Pathology laboratory (TRIP), and the Biobank, supported by the UW Department of Pathology and Laboratory Medicine, UWCCC (P30 CA014520) and the Office of The Director-NIH (S10OD023526). We acknowledge the donation of Sprague-Dawley rat skins from Dr. Kumar (Department of Comparative Biosciences, UW).

## Abbreviations

dWAT: dermal white adipose tissue
scWAT: subcutaneous white adipose tissue
BAT: brown adipose tissue
SAF: skin-associated fat
iWAT: inguinal (mouse subcutaneous) white adipose tissue
vWAT: visceral white adipose tissue
MRI: magnetic resonance imaging

## TRANSLATIONAL PERSPECTIVE

- Implicates a large skin-associated adipose depot (10-24 kG) as a potential source of heat production in humans; thus an alternative to the better-studied brown adipose tissue, an adipose depot known to be dedicated to heat production
- Documents the determinants of skin associated depots in human and mouse (sex, age, obesity, diet, cold exposure)
- Notes that the combination of dWAT with the subcutaneous depots in mouse may be the functional homolog of human skin-associated fat, which is important for modeling the physiological regulation of heat loss through mammalian skins

## Supporting Information

**Fig. S1. (**supplemental to Fig. 1). Female specific pattern of skin-associated fat accumulation.

**Fig. S2**. (supplemental to Fig. 2E, Fig. 3). Response of mouse dWAT to obesogenesis and dietary restriction **Fig. S3**. (supplemental to Fig. 5, 7). Evaluation of the specificity of immunofluorescent stains, for HSL (HSL stim/pS^660^ and HSL inh/pS^565^) and fatty acid synthase enzyme (FASN).

**Fig. S4**. (supplemental to Fig. 5). Additional markers of β-adrenergic responsivity.

**Fig. S5**. (supplemental to Fig. 5C). Lipolytic response of mouse iWAT, but not dWAT, in response to 4^0^C housing.

**Fig. S6**. (supplemental to Fig. 7). Additional responses of human skin-associated fat to the β-adrenergic agonist, mirabegron (UCP1, pHSLstim and pHSLinh).

**Fig. S7**. (supplemental to Fig. 7). Leg skin-associated fat also becomes lipolytic upon β-adrenergic stimulation.

## REFERENCES

Alexander CM, Kasza I, Yen CL, Reeder SB, Hernando D, Gallo RL, Jahoda CA, Horsley V & MacDougald OA. (2015). Dermal white adipose tissue: a new component of the thermogenic response. J Lipid Res 56, 2061–2069.

Bartelt A & Heeren J. (2014). Adipose tissue browning and metabolic health. Nat Rev Endocrinol 10, 24–36.

Cannon B & Nedergaard J. (2009). Thermogenesis challenges the adipostat hypothesis for body-weight control. Proc Nutr Soc 68, 401–407.

Cappellano G, Morandi EM, Rainer J, Grubwieser P, Heinz K, Wolfram D, Bernhard D, Lobenwein S, Pierer G & Ploner C. (2018). Human Macrophages Preferentially Infiltrate the Superficial Adipose Tissue. Int J Mol Sci 19.

Chen KY, Cypess AM, Laughlin MR, Haft CR, Hu HH, Bredella MA, Enerback S, Kinahan PE, Lichtenbelt W, Lin FI, Sunderland JJ, Virtanen KA & Wahl RL. (2016). Brown Adipose Reporting Criteria in Imaging STudies (BARCIST 1.0): Recommendations for Standardized FDG-PET/CT Experiments in Humans. Cell Metab 24, 210–222.

Chondronikola M & Sidossis LS. (2019). Brown and beige fat: From molecules to physiology. Biochim Biophys Acta Mol Cell Biol Lipids 1864, 91–103.

Chouchani ET, Kazak L & Spiegelman BM. (2019). New Advances in Adaptive Thermogenesis: UCP1 and Beyond. Cell Metab 29, 27–37.

Driskell RR, Jahoda CA, Chuong CM, Watt FM & Horsley V. (2014). Defining dermal adipose tissue. Exp Dermatol 23, 629–631.

Elattar S & Satyanarayana A. (2015). Can Brown Fat Win the Battle Against White Fat? J Cell Physiol 230, 2311–2317.

Enevoldsen LH, Simonsen L, Stallknecht B, Galbo H & Bulow J. (2001). In vivo human lipolytic activity in preperitoneal and subdivisions of subcutaneous abdominal adipose tissue. Am J Physiol Endocrinol Metab 281, E1110–1114.

Fabbiano S, Suarez-Zamorano N, Rigo D, Veyrat-Durebex C, Stevanovic Dokic A, Colin DJ & Trajkovski M. (2016). Caloric Restriction Leads to Browning of White Adipose Tissue through Type 2 Immune Signaling. Cell Metab 24, 434–446.

Finlin BS, Confides AL, Zhu B, Boulanger MC, Memetimin H, Taylor KW, Johnson ZR, Westgate PM, Dupont-Versteegden EE & Kern PA. (2019). Adipose Tissue Mast Cells Promote Human Adipose Beiging in Response to Cold. Sci Rep 9, 8658.

Finlin BS, Memetimin H, Confides AL, Kasza I, Zhu B, Vekaria HJ, Harfmann B, Jones KA, Johnson ZR, Westgate PM, Alexander CM, Sullivan PG, Dupont-Versteegden EE & Kern PA. (2018). Human adipose beiging in response to cold and mirabegron. JCI Insight 3.

Finlin BS, Zhu B, Confides AL, Westgate PM, Harfmann BD, Dupont-Versteegden EE & Kern PA. (2017). Mast Cells Promote Seasonal White Adipose Beiging in Humans. Diabetes 66, 1237–1246.

Fruhbeck G, Mendez-Gimenez L, Fernandez-Formoso JA, Fernandez S & Rodriguez A. (2014). Regulation of adipocyte lipolysis. Nutr Res Rev 27, 63–93.

Giles DA, Moreno-Fernandez ME, Stankiewicz TE, Graspeuntner S, Cappelletti M, Wu D, Mukherjee R, Chan CC, Lawson MJ, Klarquist J, Sunderhauf A, Softic S, Kahn CR, Stemmer K, Iwakura Y, Aronow BJ, Karns R, Steinbrecher KA, Karp CL, Sheridan R, Shanmukhappa SK, Reynaud D, Haslam DB, Sina C, Rupp J, Hogan SP & Divanovic S. (2017). Thermoneutral housing exacerbates nonalcoholic fatty liver disease in mice and allows for sex-independent disease modeling. Nat Med 23, 829–838.

Grove KL, Fried SK, Greenberg AS, Xiao XQ & Clegg DJ. (2010). A microarray analysis of sexual dimorphism of adipose tissues in high-fat-diet-induced obese mice. Int J Obes (Lond) 34, 989–1000.

Hanssen MJ, van der Lans AA, Brans B, Hoeks J, Jardon KM, Schaart G, Mottaghy FM, Schrauwen P & van Marken Lichtenbelt WD. (2016). Short-term Cold Acclimation Recruits Brown Adipose Tissue in Obese Humans. Diabetes 65, 1179–1189.

Harms M & Seale P. (2013). Brown and beige fat: development, function and therapeutic potential. Nature medicine 19, 1252–1263.

Ikeda K, Kang Q, Yoneshiro T, Camporez JP, Maki H, Homma M, Shinoda K, Chen Y, Lu X, Maretich P, Tajima K, Ajuwon KM, Soga T & Kajimura S. (2017). UCP1-independent signaling involving SERCA2b-mediated calcium cycling regulates beige fat thermogenesis and systemic glucose homeostasis. Nat Med 23, 1454–1465.

Kajimura S, Spiegelman BM & Seale P. (2015). Brown and Beige Fat: Physiological Roles beyond Heat Generation. Cell Metab 22, 546–559.

Karastergiou K & Fried SK. (2013). Multiple adipose depots increase cardiovascular risk via local and systemic effects. Curr Atheroscler Rep 15, 361.

Karastergiou K & Fried SK. (2017). Cellular Mechanisms Driving Sex Differences in Adipose Tissue Biology and Body Shape in Humans and Mouse Models. Adv Exp Med Biol 1043, 29–51.

Karastergiou K, Smith SR, Greenberg AS & Fried SK. (2012). Sex differences in human adipose tissues - the biology of pear shape. Biol Sex Differ 3, 13.

Kasza I, Adler D, Nelson DW, Eric Yen CL, Dumas S, Ntambi JM, MacDougald OA, Hernando D, Porter WP, Best FA & Alexander CM. (2019). Evaporative cooling provides a major metabolic energy sink. Mol Metab 27, 47–61.

Kasza I, Hernando D, Roldan-Alzate A, Alexander CM & Reeder SB. (2016). Thermogenic profiling using magnetic resonance imaging of dermal and other adipose tissues. JCI Insight 1, e87146.

Kasza I, Suh Y, Wollny D, Clark RJ, Roopra A, Colman RJ, MacDougald OA, Shedd TA, Nelson DW, Yen MI, Yen CL & Alexander CM. (2014). Syndecan-1 is required to maintain intradermal fat and prevent cold stress. PLoS Genet 10, e1004514.

Kazak L, Rahbani JF, Samborska B, Lu GZ, Jedrychowski MP, Lajoie M, Zhang S, Ramsay LC, Dou FY, Tenen D, Chouchani ET, Dzeja P, Watson IR, Tsai L, Rosen ED & Spiegelman BM. (2019). Ablation of adipocyte creatine transport impairs thermogenesis and causes diet-induced obesity. Nat Metab 1, 360–370.

Keipert S & Jastroch M. (2014). Brite/beige fat and UCP1 - is it thermogenesis? Biochim Biophys Acta 1837, 1075–1082.

Keipert S, Lutter D, Schroeder BO, Brandt D, Stahlman M, Schwarzmayr T, Graf E, Fuchs H, de Angelis MH, Tschop MH, Rozman J & Jastroch M. (2020). Endogenous FGF21-signaling controls paradoxical obesity resistance of UCP1-deficient mice. Nat Commun 11, 624.

Kelley DE, Thaete FL, Troost F, Huwe T & Goodpaster BH. (2000). Subdivisions of subcutaneous abdominal adipose tissue and insulin resistance. American journal of physiology Endocrinology and metabolism 278, E941–948.

Kern PA, Finlin BS, Zhu B, Rasouli N, McGehee RE, Jr., Westgate PM & Dupont-Versteegden EE. (2014). The effects of temperature and seasons on subcutaneous white adipose tissue in humans: evidence for thermogenic gene induction. J Clin Endocrinol Metab 99, E2772–2779.

Kershaw EE & Flier JS. (2004). Adipose tissue as an endocrine organ. J Clin Endocrinol Metab 89, 2548–2556.

Kruglikov IL & Scherer PE. (2016a). Dermal adipocytes and hair cycling: is spatial heterogeneity a characteristic feature of the dermal adipose tissue depot? Exp Dermatol 25, 258–262.

Kruglikov IL & Scherer PE. (2016b). Dermal Adipocytes: From Irrelevance to Metabolic Targets? Trends Endocrinol Metab 27, 1–10.

Kruse V, Neess D & Faergeman NJ. (2017). The Significance of Epidermal Lipid Metabolism in Whole-Body Physiology. Trends Endocrinol Metab 28, 669–683.

Kuhn JP, Meffert P, Heske C, Kromrey ML, Schmidt CO, Mensel B, Volzke H, Lerch MM, Hernando D, Mayerle J & Reeder SB. (2017). Prevalence of Fatty Liver Disease and Hepatic Iron Overload in a Northeastern German Population by Using Quantitative MR Imaging. Radiology 284, 706–716.

Langin D, Dicker A, Tavernier G, Hoffstedt J, Mairal A, Ryden M, Arner E, Sicard A, Jenkins CM, Viguerie N, van Harmelen V, Gross RW, Holm C & Arner P. (2005). Adipocyte lipases and defect of lipolysis in human obesity. Diabetes 54, 3190–3197.

Lee MJ, W. Y & Fried SK. (2013). Adipose tissue heterogeneity: implication of depot differences in adipose tissue for obesity complications. Mol Aspects Med 34, 1–11.

Liu D, Bordicchia M, Zhang C, Fang H, Wei W, Li JL, Guilherme A, Guntur K, Czech MP & Collins S. (2016). Activation of mTORC1 is essential for beta-adrenergic stimulation of adipose browning. J Clin Invest 126, 1704–1716.

Loh NY, Neville MJ, Marinou K, Hardcastle SA, Fielding BA, Duncan EL, McCarthy MI, Tobias JH, Gregson CL, Karpe F & Christodoulides C. (2015). LRP5 Regulates Human Body Fat Distribution by Modulating Adipose Progenitor Biology in a Dose- and Depot-Specific Fashion. Cell metabolism 21, 262–272.

Lu Y, Day FR, Gustafsson S, Buchkovich ML, Na J, Bataille V, Cousminer DL, Dastani Z, Drong AW, Esko T, Evans DM, Falchi M, Feitosa MF, Ferreira T, Hedman AK, Haring R, Hysi PG, Iles MM, Justice AE, Kanoni S, Lagou V, Li R, Li X, Locke A, Lu C, Magi R, Perry JR, Pers TH, Qi Q, Sanna M, Schmidt EM, Scott WR, Shungin D, Teumer A, Vinkhuyzen AA, Walker RW, Westra HJ, Zhang M, Zhang W, Zhao JH, Zhu Z, Afzal U, Ahluwalia TS, Bakker SJ, Bellis C, Bonnefond A, Borodulin K, Buchman AS, Cederholm T, Choh AC, Choi HJ, Curran JE, de Groot LC, De Jager PL, Dhonukshe-Rutten RA, Enneman AW, Eury E, Evans DS, Forsen T, Friedrich N, Fumeron F, Garcia ME, Gartner S, Han BG, Havulinna AS, Hayward C, Hernandez D, Hillege H, Ittermann T, Kent JW, Kolcic I, Laatikainen T, Lahti J, Mateo Leach I, Lee CG, Lee JY, Liu T, Liu Y, Lobbens S, Loh M, Lyytikainen LP, Medina-Gomez C, Michaelsson K, Nalls MA, Nielson CM, Oozageer L, Pascoe L, Paternoster L, Polasek O, Ripatti S, Sarzynski MA, Shin CS, Narancic NS, Spira D, Srikanth P, Steinhagen-Thiessen E, Sung YJ, Swart KM, Taittonen L, Tanaka T, Tikkanen E, van der Velde N, van Schoor NM, Verweij N, Wright AF, Yu L, Zmuda JM, Eklund N, Forrester T, Grarup N, Jackson AU, Kristiansson K, Kuulasmaa T, Kuusisto J, Lichtner P, Luan J, Mahajan A, Mannisto S, Palmer CD, Ried JS, Scott RA, Stancakova A, Wagner PJ, Demirkan A, Doring A, Gudnason V, Kiel DP, Kuhnel B, Mangino M, McKnight B, Menni C, O’Connell JR, Oostra BA, Shuldiner AR, Song K, Vandenput L, van Duijn CM, Vollenweider P, White CC, Boehnke M, Boettcher Y, Cooper RS, Forouhi NG, Gieger C, Grallert H, Hingorani A, Jorgensen T, Jousilahti P, Kivimaki M, Kumari M, Laakso M, Langenberg C, Linneberg A, Luke A, McKenzie CA, Palotie A, Pedersen O, Peters A, Strauch K, Tayo BO, Wareham NJ, Bennett DA, Bertram L, Blangero J, Bluher M, Bouchard C, Campbell H, Cho NH, Cummings SR, Czerwinski SA, Demuth I, Eckardt R, Eriksson JG, Ferrucci L, Franco OH, Froguel P, Gansevoort RT, Hansen T, Harris TB, Hastie N, Heliovaara M, Hofman A, Jordan JM, Jula A, Kahonen M, Kajantie E, Knekt PB, Koskinen S, Kovacs P, Lehtimaki T, Lind L, Liu Y, Orwoll ES, Osmond C, Perola M, Perusse L, Raitakari OT, Rankinen T, Rao DC, Rice TK, Rivadeneira F, Rudan I, Salomaa V, Sorensen TI, Stumvoll M, Tonjes A, Towne B, Tranah GJ, Tremblay A, Uitterlinden AG, van der Harst P, Vartiainen E, Viikari JS, Vitart V, Vohl MC, Volzke H, Walker M, Wallaschofski H, Wild S, Wilson JF, Yengo L, Bishop DT, Borecki IB, Chambers JC, Cupples LA, Dehghan A, Deloukas P, Fatemifar G, Fox C, Furey TS, Franke L, Han J, Hunter DJ, Karjalainen J, Karpe F, Kaplan RC, Kooner JS, McCarthy MI, Murabito JM, Morris AP, Bishop JA, North KE, Ohlsson C, Ong KK, Prokopenko I, Richards JB, Schadt EE, Spector TD, Widen E, Willer CJ, Yang J, Ingelsson E, Mohlke KL, Hirschhorn JN, Pospisilik JA, Zillikens MC, Lindgren C, Kilpelainen TO & Loos RJ. (2016). New loci for body fat percentage reveal link between adiposity and cardiometabolic disease risk. Nat Commun 7, 10495.

Manolopoulos KN, Karpe F & Frayn KN. (2010). Gluteofemoral body fat as a determinant of metabolic health. Int J Obes (Lond) 34, 949–959.

Mills EL, Pierce KA, Jedrychowski MP, Garrity R, Winther S, Vidoni S, Yoneshiro T, Spinelli JB, Lu GZ, Kazak L, Banks AS, Haigis MC, Kajimura S, Murphy MP, Gygi SP, Clish CB & Chouchani ET. (2018). Accumulation of succinate controls activation of adipose tissue thermogenesis. Nature 560, 102–106.

Mottillo EP, Balasubramanian P, Lee YH, Weng C, Kershaw EE & Granneman JG. (2014). Coupling of lipolysis and de novo lipogenesis in brown, beige, and white adipose tissues during chronic beta3-adrenergic receptor activation. J Lipid Res 55, 2276–2286.

Neess D, Bek S, Bloksgaard M, Marcher AB, Faergeman NJ & Mandrup S. (2013). Delayed hepatic adaptation to weaning in ACBP-/-mice is caused by disruption of the epidermal barrier. Cell Rep 5, 1403–1412.

Neess D, Kruse V, Marcher AB, Waede MR, Vistisen J, Moller PM, Petersen R, Brewer JR, Ma T, Colleluori G, Severi I, Cinti S, Gerhart-Hines Z, Mandrup S & Faergeman NJ. (2020). Epidermal Acyl-CoA-binding protein is indispensable for systemic energy homeostasis. Mol Metab 44, 101144.

Nicu C, Pople J, Bonsell L, Bhogal R, Ansell DM & Paus R. (2018). A guide to studying human dermal adipocytes in situ. Exp Dermatol 27, 589–602.

Ogasawara J, Izawa T, Sakurai T, Sakurai T, Shirato K, Ishibashi Y, Ishida H, Ohno H & Kizaki T. (2015). The Molecular Mechanism Underlying Continuous Exercise Training-Induced Adaptive Changes of Lipolysis in White Adipose Cells. J Obes 2015, 473430.

Perez-Chirinos Buxade C, Sola-Perez T, Castizo-Olier J, Carrasco-Marginet M, Roy A, Marfell-Jones M & Irurtia A. (2018). Assessing subcutaneous adipose tissue by simple and portable field instruments: Skinfolds versus A-mode ultrasound measurements. PLoS One 13, e0205226.

Pinnick KE, Nicholson G, Manolopoulos KN, McQuaid SE, Valet P, Frayn KN, Denton N, Min JL, Zondervan KT, Fleckner J, Mol PC, McCarthy MI, Holmes CC & Karpe F. (2014). Distinct developmental profile of lower-body adipose tissue defines resistance against obesity-associated metabolic complications. Diabetes 63, 3785–3797.

Qiao G, Chen M, Bucsek MJ, Repasky EA & Hylander BL. (2018). Adrenergic Signaling: A Targetable Checkpoint Limiting Development of the Antitumor Immune Response. Front Immunol 9, 164.

Rivera-Gonzalez GC, Shook BA, Andrae J, Holtrup B, Bollag K, Betsholtz C, Rodeheffer MS & Horsley V. (2016). Skin Adipocyte Stem Cell Self-Renewal Is Regulated by a PDGFA/AKT-Signaling Axis. Cell Stem Cell 19, 738–751.

Sampath H, Flowers MT, Liu X, Paton CM, Sullivan R, Chu K, Zhao M & Ntambi JM. (2009). Skin-specific deletion of stearoyl-CoA desaturase-1 alters skin lipid composition and protects mice from high fat diet-induced obesity. The Journal of biological chemistry 284, 19961–19973.

Sampath H & Ntambi JM. (2014). Role of stearoyl-CoA desaturase-1 in skin integrity and whole body energy balance. J Biol Chem 289, 2482–2488.

Sbarbati A, Accorsi D, Benati D, Marchetti L, Orsini G, Rigotti G & Panettiere P. (2010). Subcutaneous adipose tissue classification. Eur J Histochem 54, e48.

Scheller EL, Khandaker S, Learman BS, Cawthorn WP, Anderson LM, Pham HA, Robles H, Wang Z, Li Z, Parlee SD, Simon BR, Mori H, Bree AJ, Craft CS & MacDougald OA. (2019). Bone marrow adipocytes resist lipolysis and remodeling in response to beta-adrenergic stimulation. Bone 118, 32–41.

Shih MY, Kane MA, Zhou P, Yen CL, Streeper RS, Napoli JL & Farese RV, Jr. (2009). Retinol Esterification by DGAT1 Is Essential for Retinoid Homeostasis in Murine Skin. J Biol Chem 284, 4292–4299.

Smith GI, Mittendorfer B & Klein S. (2019). Metabolically healthy obesity: facts and fantasies. J Clin Invest 129, 3978–3989.

Smith SR, Lovejoy JC, Greenway F, Ryan D, deJonge L, de la Bretonne J, Volafova J & Bray GA. (2001). Contributions of total body fat, abdominal subcutaneous adipose tissue compartments, and visceral adipose tissue to the metabolic complications of obesity. Metabolism 50, 425–435.

Speakman JR. (2013). Measuring energy metabolism in the mouse - theoretical, practical, and analytical considerations. Front Physiol 4, 34.

Storchle P, Muller W, Sengeis M, Lackner S, Holasek S & Furhapter-Rieger A. (2018). Measurement of mean subcutaneous fat thickness: eight standardised ultrasound sites compared to 216 randomly selected sites. Sci Rep 8, 16268.

Tian XY, Ganeshan K, Hong C, Nguyen KD, Qiu Y, Kim J, Tangirala RK, Tonotonoz P & Chawla A. (2016). Thermoneutral Housing Accelerates Metabolic Inflammation to Potentiate Atherosclerosis but Not Insulin Resistance. Cell Metab 23, 165–178.

Tschop MH, Speakman JR, Arch JR, Auwerx J, Bruning JC, Chan L, Eckel RH, Farese RV, Jr., Galgani JE, Hambly C, Herman MA, Horvath TL, Kahn BB, Kozma SC, Maratos-Flier E, Muller TD, Munzberg H, Pfluger PT, Plum L, Reitman ML, Rahmouni K, Shulman GI, Thomas G, Kahn CR & Ravussin E. (2012). A guide to analysis of mouse energy metabolism. Nature methods 9, 57–63.

van der Lans AA, Hoeks J, Brans B, Vijgen GH, Visser MG, Vosselman MJ, Hansen J, Jorgensen JA, Wu J, Mottaghy FM, Schrauwen P & van Marken Lichtenbelt WD. (2013). Cold acclimation recruits human brown fat and increases nonshivering thermogenesis. J Clin Invest 123, 3395–3403.

van Marken Lichtenbelt WD, Vanhommerig JW, Smulders NM, Drossaerts JM, Kemerink GJ, Bouvy ND, Schrauwen P & Teule GJ. (2009). Cold-activated brown adipose tissue in healthy men. The New England journal of medicine 360, 1500–1508.

Verboven K, Wouters K, Gaens K, Hansen D, Bijnen M, Wetzels S, Stehouwer CD, Goossens GH, Schalkwijk CG, Blaak EE & Jocken JW. (2018). Abdominal subcutaneous and visceral adipocyte size, lipolysis and inflammation relate to insulin resistance in male obese humans. Sci Rep 8, 4677.

Villarroya F & Giralt M. (2015). The Beneficial Effects of Brown Fat Transplantation: Further Evidence of an Endocrine Role of Brown Adipose Tissue. Endocrinology 156, 2368–2370.

Volzke H, Alte D, Schmidt CO, Radke D, Lorbeer R, Friedrich N, Aumann N, Lau K, Piontek M, Born G, Havemann C, Ittermann T, Schipf S, Haring R, Baumeister SE, Wallaschofski H, Nauck M, Frick S, Arnold A, Junger M, Mayerle J, Kraft M, Lerch MM, Dorr M, Reffelmann T, Empen K, Felix SB, Obst A, Koch B, Glaser S, Ewert R, Fietze I, Penzel T, Doren M, Rathmann W, Haerting J, Hannemann M, Ropcke J, Schminke U, Jurgens C, Tost F, Rettig R, Kors JA, Ungerer S, Hegenscheid K, Kuhn JP, Kuhn J, Hosten N, Puls R, Henke J, Gloger O, Teumer A, Homuth G, Volker U, Schwahn C, Holtfreter B, Polzer I, Kohlmann T, Grabe HJ, Rosskopf D, Kroemer HK, Kocher T, Biffar R, John U & Hoffmann W. (2011). Cohort profile: the study of health in Pomerania. Int J Epidemiol 40, 294–307.

Wajchenberg BL. (2000). Subcutaneous and visceral adipose tissue: their relation to the metabolic syndrome. Endocr Rev 21, 697–738.

Walker GE, Marzullo P, Ricotti R, Bona G & Prodam F. (2014). The pathophysiology of abdominal adipose tissue depots in health and disease. Horm Mol Biol Clin Investig 19, 57–74.

Wang GX, Zhao XY & Lin JD. (2015). The brown fat secretome: metabolic functions beyond thermogenesis Trends Endocrinol Metab 26, 231–237.

Wu J, Bostrom P, Sparks LM, Ye L, Choi JH, Giang AH, Khandekar M, Virtanen KA, Nuutila P, Schaart G, Huang K, Tu H, van Marken Lichtenbelt WD, Hoeks J, Enerback S, Schrauwen P & Spiegelman BM. (2012). Beige adipocytes are a distinct type of thermogenic fat cell in mouse and human. Cell 150, 366–376.

Yoneshiro T, Aita S, Matsushita M, Kayahara T, Kameya T, Kawai Y, Iwanaga T & Saito M. (2013). Recruited brown adipose tissue as an antiobesity agent in humans. J Clin Invest 123, 3404–3408.

Yoneshiro T, Matsushita M, Nakae S, Kameya T, Sugie H, Tanaka S & Saito M. (2016). Brown adipose tissue is involved in the seasonal variation of cold-induced thermogenesis in humans. Am J Physiol Regul Integr Comp Physiol, ajpregu 00057 02015.

Zhang Z, Shao M, Hepler C, Zi Z, Zhao S, An YA, Zhu Y, Ghaben AL, Wang MY, Li N, Onodera T, Joffin N, Crewe C, Zhu Q, Vishvanath L, Kumar A, Xing C, Wang QA, Gautron L, Deng Y, Gordillo R, Kruglikov I, Kusminski CM, Gupta RK & Scherer PE. (2019). Dermal adipose tissue has high plasticity and undergoes reversible dedifferentiation in mice. J Clin Invest 129, 5327–5342.

Zwick RK, Guerrero-Juarez CF, Horsley V & Plikus MV. (2018). Anatomical, Physiological, and Functional Diversity of Adipose Tissue. Cell Metab 27, 68–83.

