## Supplemental Figures for "Contrasting recruitment of skin-associated adipose depots during cold challenge of mouse and human" (from another version of this preprint: version 1, 2020-09-18)

Fig. S1

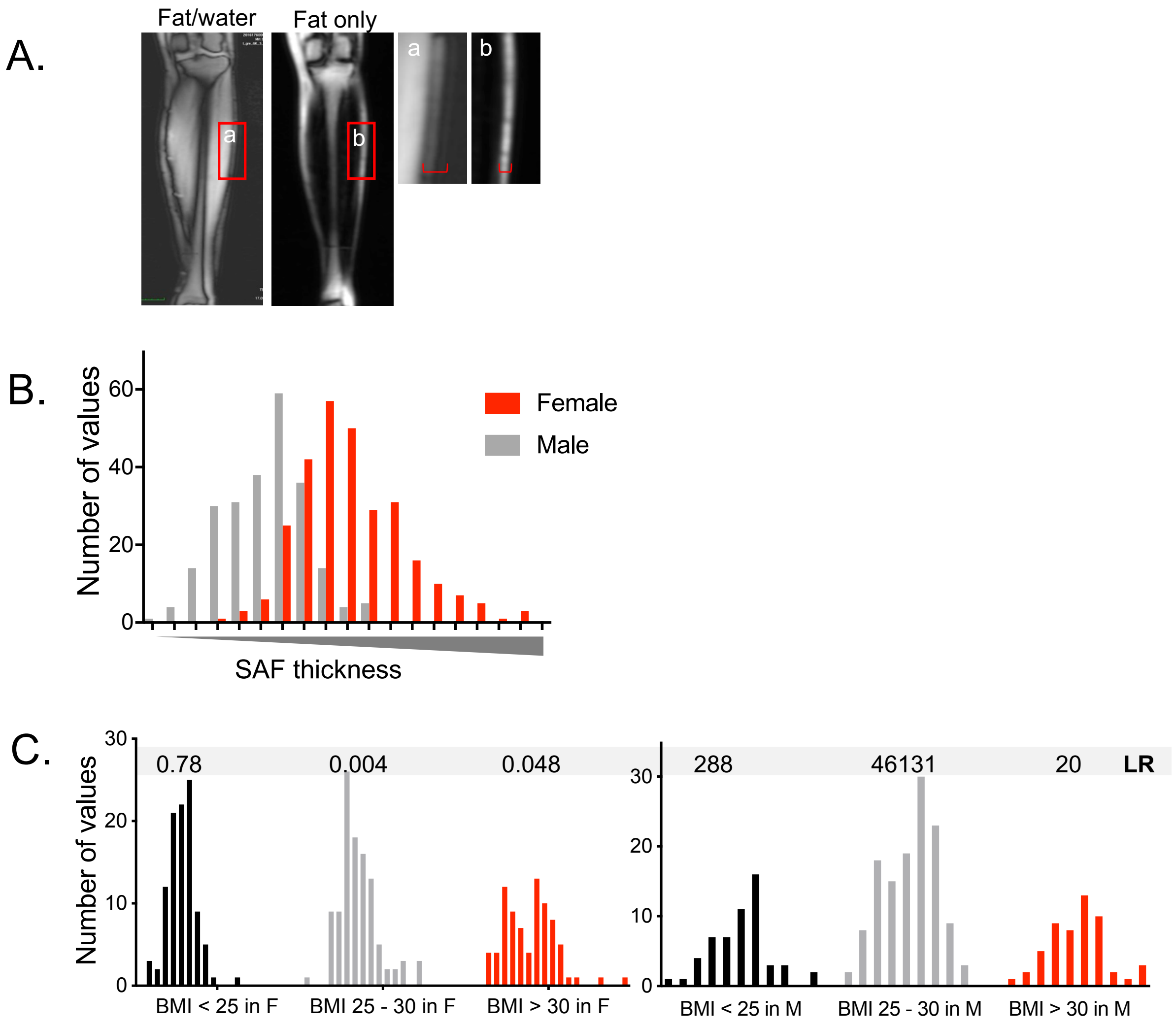


**Fig. S1.** (for Fig. 1). **Female specific pattern of skin-associated fat accumulation. A.** Comparison of images collected using a standard fat/water protocol versus fat only. **B.** SAF thickness was binned into groups for all men and women, to illustrate the relative overlap and shape of each curve. **C.** The distribution of SAF thickness for lean, over-weight and obese women is compared with the same parameters in men; the likelihood ratio (LR) for normal versus lognormal distribution is calculated, generating the conclusion that the SAF thickness for men follows a normal distribution, but not for women (BMI<25, p=0.001; BMI 25-30, p<0.0001; BMI.30, p-0.0063; D'Agostino and Pearson test).

Fig. S2

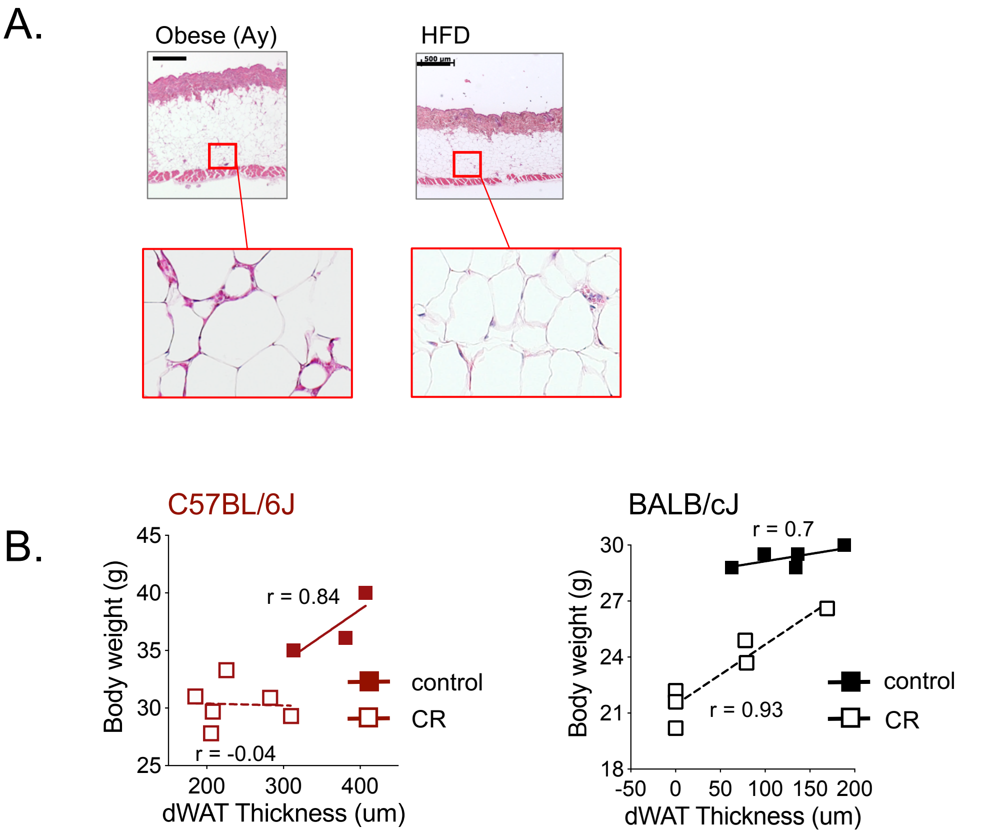


**Fig. S2.** (Fig. 2D, 3). **Response of dWAT to body-wide fat accumulation. A.** *dWAT is quite resistant to crowning/inflammatory macrophage influx.* Crown like structures can be identified from H&E stained sections of skins of genetically obese ($A^y$) mice (>30 weeks old) but not in mice fed a high fat diet for 26 weeks as follows: diet#TD.88137 for 12 weeks, then an amino acid defined high fat diet, TD.160186 for 14 weeks, both from Envigo, Madison WI. Complete design of this study is detailed in (68). Scale bars=500 µm. **B.** *Calorie restricted mice show strong correlation of overall body weight and dWAT thickness.* Body weight and dWAT thickness are correlated for both control and calorie restricted BALB/cJ mice. The leanest calorie restricted BALB/cJ mice show negligible dWAT levels, whereas C57BL/6J mice protect at least 200 µm thick dWAT. Pearson correlation coefficients were calculated

Fig. S3

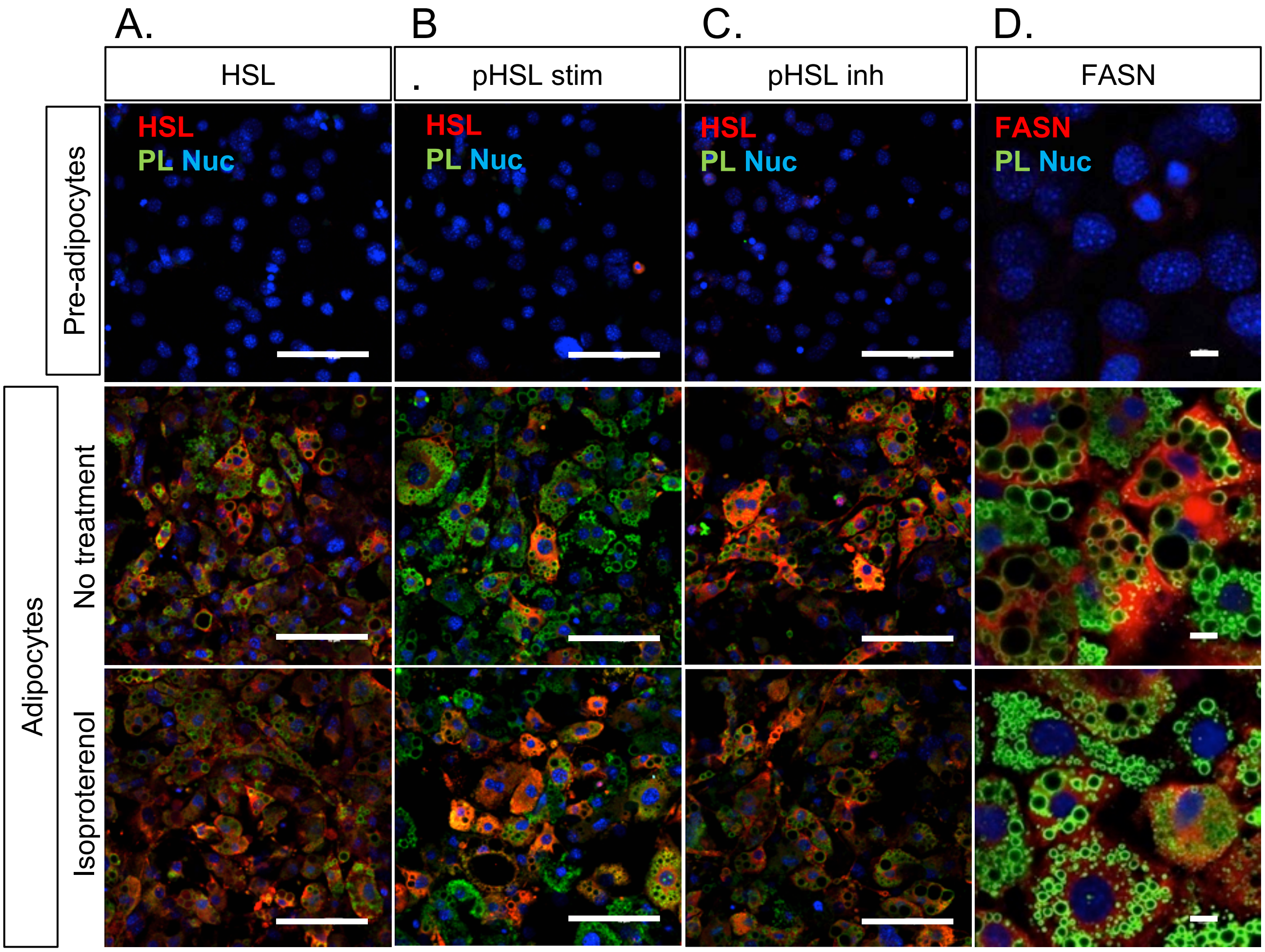


**Fig. S3.** (Fig. 5, 7). **Evaluation of the specificity of immunofluorescent stains, for HSL (HSL stim/pS$^{660}$ and HSL inh/pS$^{565}$) and fatty acid synthase enzyme (FASN).** 3T3 L1 cells, either undifferentiated or after treatment to induce adipogenesis, were stained with antibodies for HSL or FASN (red), in combination with perilipin (green) and DAPI (blue; nuc; nuclei) with and without isoproterenol treatment. Total HSL staining of 3T3-L1 adipocytes was unchanged, where pHSL stim increased and pHSL inh and FASN signal were reduced. Scale bars=50 μm (A,B,C), 10 μm (D).

Fig. S4

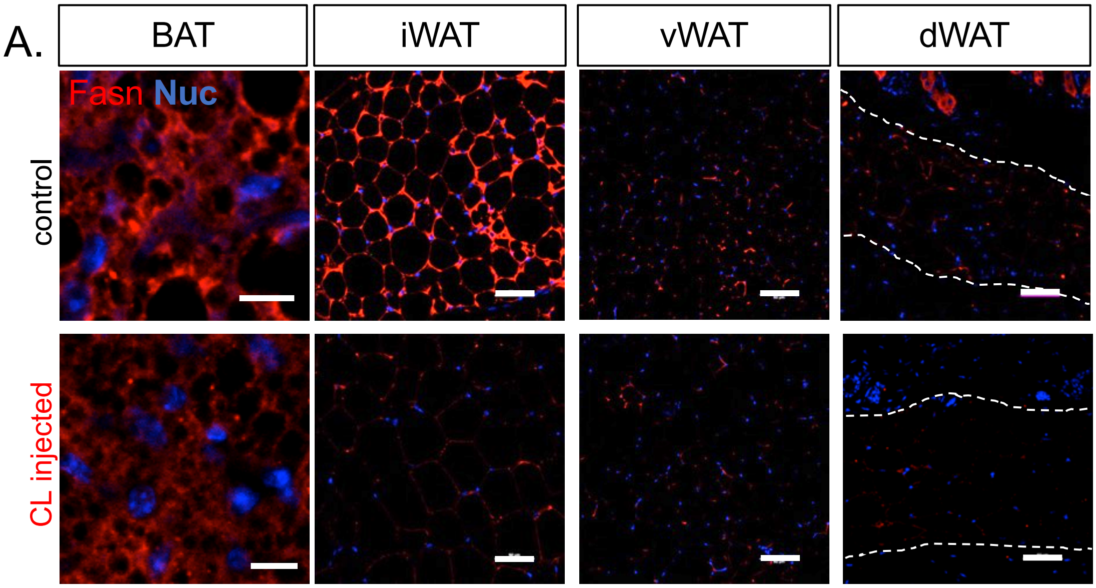


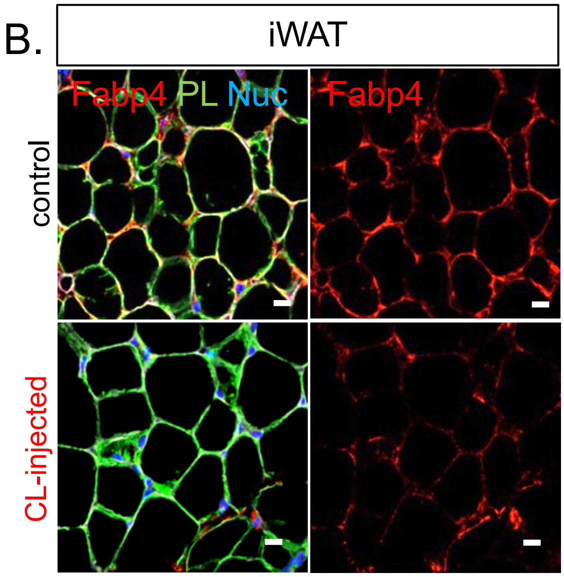


**Fig. S4.** (Fig. 5). **Additional markers of β-adrenergic responsivity. A.** *Signal for FASN is suppressed by β-adrenergic induction in beige/brown depots, but not dWAT/vWAT.* The same panel of tissues shown in Fig. 5 (BAT, iWAT, vWAT and dWAT), from CL-injected mice or controls, were stained for FASN to illustrate that the signal for this lipogenic enzyme is suppressed in thermogenic depots (iWAT and BAT) but no signal is observed in either vWAT or dWAT (induced or not). Scale bars=20 µm (BAT) or 100 µm (iWAT, vWAT, dWAT). **B.** *Signal for Fabp4 is suppressed by β-adrenergic induction in beige fat, but not dWAT/vWAT.* Similar stains of FABP4 show that this lipogenic activity is likewise suppressed in iWAT and shows little signal in either vWAT or dWAT. Scale bars=20 µm (iWAT) or 100 µm (vWAT, dWAT).

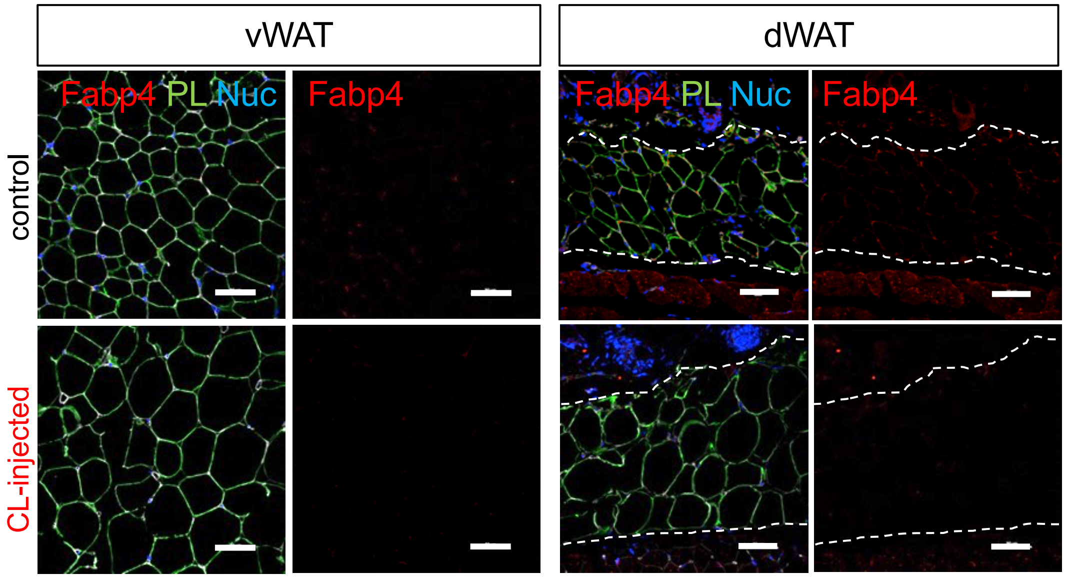

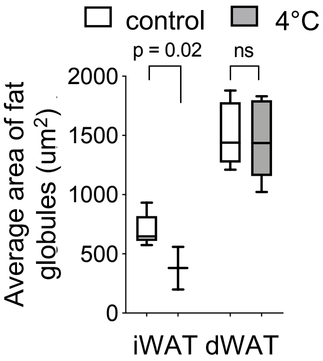


**Fig. S5.** (Fig. 5C). **Lipolytic response of mouse iWAT, but not dWAT, in response to $4^0$C housing.** The average area of adipocyte-associated fat globules was measured as described for Fig. 5C, after transfer of mice from room temperature (control) to $4^0$C for 1 hour. Data were analyzed using an unpaired 2-tailed *t* test.

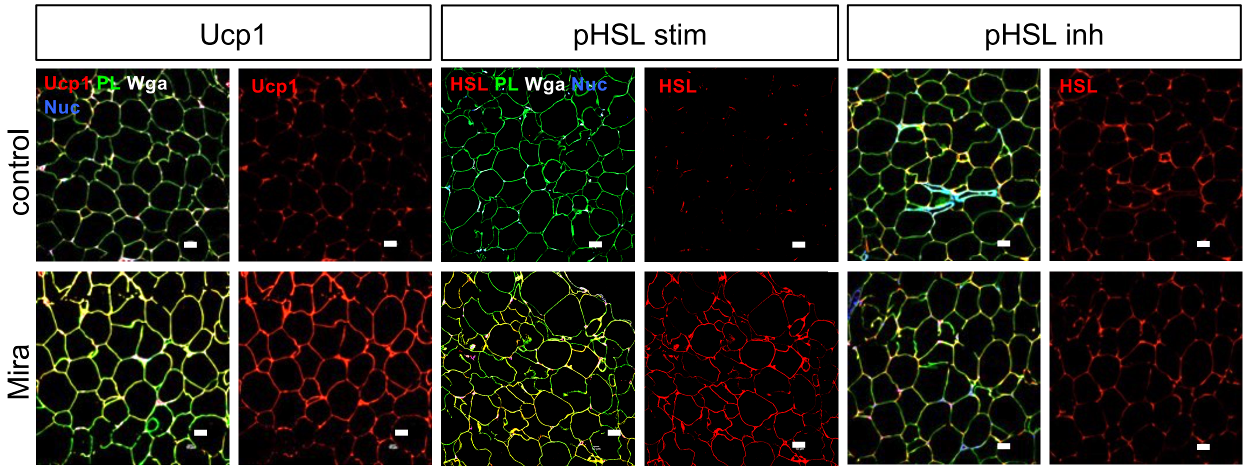


**Fig. S6.** (Fig. 7). **Additional responses of human skin-associated fat to the β-adrenergic agonist, mirabegron.** In abdominal SAF from obese subjects, administered the β3-adrenergic agent mirabegron daily, immunostains for UCP1 (red) and regulatory modifications of HSL (red) (pHSL stim and pHSL inh) in combination with perilipin (green) and DAPI (blue; nuc; nuclei) show relatively uniform activation of lipolysis throughout the entire community of adipocytes. Scale bars=20 μm.

A.

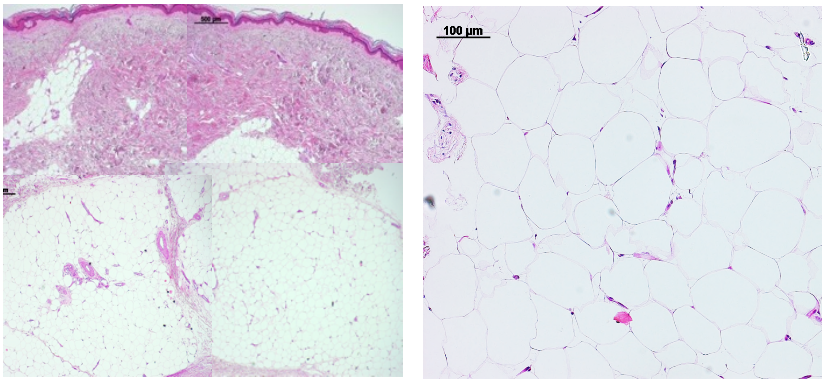


B.

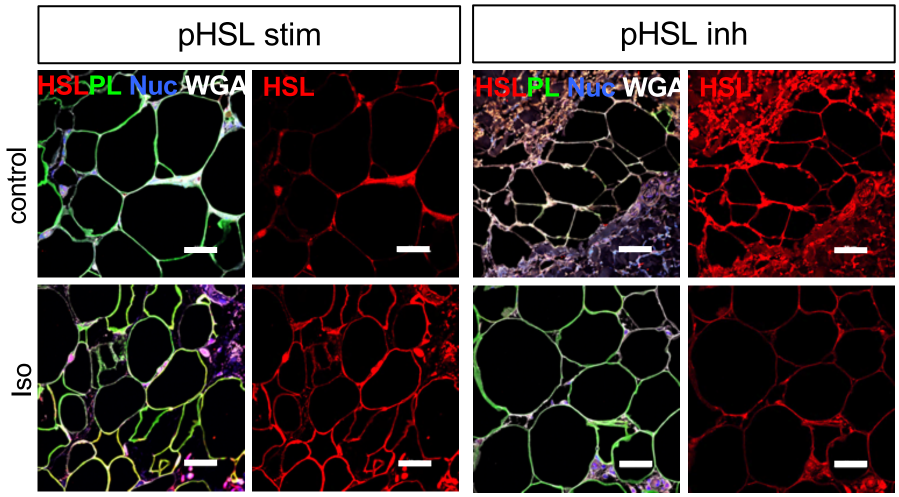


**Fig. S7.** (Fig. 7). **Leg skin-associated fat also become lipolytic upon β-adrenergic stimulation. A.** H&E sections of mid-calf-level skin show this adipose depot to be indistinguishable from skin-associated fat from other body sites. Scale bar=500 μm (left hand side); 100 μm (right hand side). **B.** Leg skin samples was treated *ex vivo*, according to Fig. 7A, to show increased staining of HSL stim (red), and reduced staining of pHSL inh (red) in combination with perilipin (green), DAPI (blue; nuc; nuclei) and wheat germ agglutinin (WGA, white, cell membrane). Scale bars=50 μm.
